# Rapid floral syndrome convergence in *Penstemon* through independent genetic variation

**DOI:** 10.64898/2026.07.21.739878

**Authors:** Benjamin W. Stone, Lucas C. Wheeler, Palmer W. Lambert, Noah H. Williams, Carolyn A. Wessinger

## Abstract

The repeated evolution of certain complex traits within a given lineage is compelling and presents an opportunity for understanding selective and genetic features that promote rapid multi-trait adaptation. The North American plant genus *Penstemon* is one such example, with numerous evolutionary shifts from ancestral bee syndrome to hummingbird syndrome flowers. Here, we traced the evolution of multi-trait floral phenotypes within a focal clade of *Penstemon* using a new whole genome phylogenomic estimate and found four independent origins of the hummingbird syndrome. We found strong evolutionary convergence in floral traits across these four origins and observed that the hummingbird syndrome assembled rapidly, without leaving a signal of stepwise modification of traits. Patterns of evolutionary correlations among floral traits in bee syndrome *Penstemon* species likely enable rapid shifts to hummingbird syndrome. Using phylogenomic tests for introgression, we found no evidence that adaptive introgression has fueled the four repeated origins of hummingbird syndrome flowers. Patterns of allele sharing were instead consistent with substantial levels of incomplete lineage sorting, suggesting repeated complex adaptation involves de novo mutation or adaptation from ancestral variation.

## Introduction

Adaptive syndromes are multi-trait adaptations that have repeatedly evolved in response to similar ecological conditions (reviewed by Losos, 2011; Sinnott-Armstrong et al., 2022; Thomson & Wilson, 2008). Multiple shifts in adaptive syndrome are common in adaptive radiations. Well-studied examples include repeated origins of freshwater morph in sticklebacks (Bell & Foster, 1994), ecomorphs in Caribbean *Anolis* lizards (Beuttell & Losos, 1999), C4 photosynthesis in grasses (Christin et al., 2013), cave-dwelling syndrome in cavefish (Dowling et al., 2002), and wing pattern mimicry in *Heliconius* butterflies (Brower, 1996). This phenomenon, while familiar, remains enigmatic. It suggests that certain lineages may be prone to a particular evolutionary shift in syndrome, despite it requiring evolutionary change to multiple component traits. These observations raise the question: why do certain adaptive syndromes evolve frequently within some lineages?

In some cases, perhaps the ancestral phenotype is an exceptional starting point that can jumpstart the step-wise assembly of the novel syndrome. In particular, ancestral traits can represent initial steps towards the novel syndrome, even if they evolved in a different ecological context (Gould & Vrba, 1982). For example, evolutionary changes to bundle sheath cell physiology unrelated to photosynthesis appears to have facilitated repeated origins of C4 photosynthesis in grasses (Burgess & Hibberd, 2015; Swift et al., 2024), and the hydrophobic skin of *Anolis* lizards seems to have facilitated repeated origins of underwater bubble-based respiration in aquatic diving species (Boccia et al., 2021).

Genetic features might also predispose a lineage to evolve a particular syndrome. These features include genetic correlations among traits that are parallel to the direction of adaptation (Schluter, 1996; Stebbins, 1974), as seen in *Anolis* where repeated trait divergence among ecomorphs aligns with patterns of genetic covariance among skeletal dimension traits (McGlothlin et al., 2018). A ready supply of adaptive alleles may also promote repeated shifts – either because relevant alleles arise frequently through new mutation due to a large mutational target size (e.g., Chan et al., 2010; Galen et al., 2015; Stoltzfus & McCandlish, 2017; Wessinger & Rausher, 2015), or because adaptive alleles are already available as standing genetic variation or shared between populations through gene flow (e.g., Enbody et al., 2023; Hooper et al., 2024; Jones et al., 2018; Short & Streisfeld, 2023). For example, repeated shifts from marine to freshwater morphs in threespine sticklebacks have involved both loss-of-function mutations that arise frequently (Chan et al., 2010) and recurrent adaptation from standing genetic variation (Colosimo et al., 2005). Given that complex trait evolution requires change in multiple component traits, adaptive introgression following hybridization events could circumvent the long waiting times expected for de novo mutations. For example, repeated shifts in both mimicry patterns and preference for these patterns in *Heliconius* butterflies have involved adaptive introgression of causal alleles (Pardo-Diaz et al., 2012; Rossi et al., 2024). Despite these accumulating examples, it remains unclear how frequently these different sources of genetic variation contribute to repeated shifts in syndrome. Whether complex adaptations involve de novo mutations, standing genetic variation, or adaptive introgression informs on whether trait evolution is constrained by mutation events, the maintenance of ancestral variation, or the opportunity for hybridization between differentiated lineages.

A striking example of convergent evolution of complex adaptation is the hummingbird pollination syndrome of western North American flora. This floral syndrome – bright red narrowly tubular flowers with exserted reproductive organs that produce copious nectar – has convergently evolved in approximately 39 plant genera, leading to at least 129 species (Abrahamczyk & Renner, 2015; Grant & Grant, 1968; Stebbins, 1989). The North American perennial wildflower genus *Penstemon* is an outlier for this pattern of convergence. *Penstemon* has experienced a rapid continental radiation into diverse ecological niches over the past 2 million years (Wolfe et al., 2021; Wolfe et al., 2006), including repeated shifts in floral pollination syndrome. Most of the approximately 290 species in the genus are bee- or wasp-pollinated and have bee syndrome flowers: flowers are blue, purple, or pale, produce small amounts of concentrated nectar, are relatively wide, and have short stamens and styles. Hummingbird-adapted flowers have evolved in at least twenty lineages (Wessinger et al., 2016; Wilson et al., 2007; Wolfe et al., 2006), and are bright red, produce large amounts of dilute nectar, have a relatively long and narrow tubular shape, and have long stamens and styles that often extend beyond the floral opening. These distinct combinations of floral traits predict pollinator visitation in *Penstemon* (Wilson et al., 2004). Evolutionary transitions to other pollination systems, such as moth, butterfly, wind, or self-pollination, are not observed or are quite rare in this genus. This remarkably widespread yet specific pattern of convergent evolution makes *Penstemon* an exceptional system to quantify evolutionary convergence and identify the source of genetic variation for these shifts.

Whether pre-existing floral features in bee-adapted penstemons promote evolutionary shifts to hummingbird pollination is currently unclear. The current conceptual model proposes a stepwise construction of the floral syndrome (Thomson & Wilson, 2008; Wilson et al., 2004; Wilson et al., 2007), where “pro-bird” adaptations to increase the frequency and efficiency of hummingbird visitation precede the evolution of “anti-bee” adaptations that deter visitation by bees after plants have specialized on bird pollinators (Castellanos et al., 2004). In particular, increased nectar production is a likely initial step that spurs shifts to hummingbird syndrome (Thomson & Wilson, 2008). This model predicts that bee-adapted species with increased nectar offerings should be close relatives to hummingbird syndrome species. Tests of a stepwise assembly using phylogenetic comparative methods have been limited by poor phylogenetic resolution that stems from the rapid radiation of the genus.

Another proposed explanation for the widespread repeated origins of hummingbird syndrome in *Penstemon* is adaptive introgression of relevant alleles following hybridization events (Stebbins, 1989). This hypothesis is plausible – hummingbirds are capable of pollinating bee syndrome *Penstemon* species (e.g., Castellanos et al., 2003) and hybrids between bee and hummingbird syndrome species can occasionally be found in sympatric populations (e.g., Crosswhite, 1965; Crump et al., 2020; de los Santos-Gómez & Ornelas, 2025; Kimball, 2008; Wilson & Valenzuela, 2002). A role for adaptive introgression would help explain why shifts to hummingbird pollination have so greatly outnumbered shifts to other pollination systems in the genus. It would further suggest that shifts to hummingbird syndrome might be limited by geographic proximity to other hummingbird syndrome penstemons.

Here, we characterized phylogenetic patterns of floral diversification in a focal clade within *Penstemon* that includes four species described as hummingbird syndrome. Using whole genome resequencing data, we estimated phylogenomic relationships within this clade. We analyzed phenotypic data to determine whether sampled species separate into distinct clusters in multivariate trait space corresponding to the classic syndromes, characterize the extent and tempo of multi-trait convergence, and examine whether hummingbird syndrome traits are assembled in a stepwise manner. In addition, we characterized patterns of phylogenomic discordance to examine whether adaptive introgression or ancestral genetic variation has served as a source of genetic variation for repeated shifts to hummingbird syndrome.

## Methods

### Study system and sampling

Our study focused on *Penstemon* section *Habroanthus* (hereafter “*Habroanthus*”), a focal clade for examining phylogenetic complexities within *Penstemon* and their contribution to convergent floral evolution. *Habroanthus* is a large western North American clade comprised of approximately 50 species (Freeman, 2019). Four species within this section are described as hummingbird syndrome and all other species are described as bee syndrome. This section has been the focus of ecological, quantitative genetic, and population genomic studies (Castellanos et al., 2003; Wessinger et al., 2014; Wessinger et al., 2023; Wessinger et al., 2018). Reference genomes have been assembled for two members of the section, *P. barbatus* (Wessinger et al., 2023) and *P. eatonii* (Jarvis et al., 2025).

We sampled 52 accessions, each from a unique population, for DNA sequencing. These accessions represent 20 *Habroanthus* species and one member of *Penstemon* section *Spectabiles* (*P. palmeri*) to serve as an outgroup. Sampling localities and voucher information for all sampled specimens can be found in Table S1. While we sampled all major lineages within *Habroanthus*, we focused our sampling on each of the four hummingbird syndrome species and closely related bee syndrome species, according to current phylogenetic estimates (Wessinger et al., 2016; Wolfe et al., 2021). This sampling was well-suited to identify the major processes contributing to genetic differentiation between lineages and to floral adaptation within the clade.

### Phenotyping

We collected floral phenotypic data for 34 accessions, measuring at least three flowers per individual (1-3 individuals per accession). For all but four of these accessions, we collected individual plants and brought them back to the plant growth facilities at University of South Carolina, where we reared them until flowering. The remaining four accessions were measured in the field for all traits except nectar production and petal spectral reflectance.

We measured floral dimension traits (tube length and width, long and short stamen lengths, and style length), nectar volume, and nectary area. We extracted and identified anthocyanidin pigments from pressed floral tissue and scored for the presence of each type using a qualitative scale. We measured petal reflectance spectra using UV-VIS spectroscopy and then estimated the detectability of each floral reflectance spectrum to bees vs. hummingbirds by comparing chromatic contrast values under a honeybee visual model vs. an average avian violet sensitive model. Further details on our phenotypic measurements are provided in the supplemental methods.

We calculated principal components from our multi-trait dataset using the R package *MorphoTools2* (Šlenker et al., 2022), excluding traits based on reflectance spectroscopy due to missing data. We log-transformed nectar volume and nectary area measurements for all analyses.

### Whole genome resequencing

We extracted DNA from fresh or silica-dried leaf tissue using a modified CTAB protocol and generated whole genome Illumina resequencing data for each sample at the Duke University Sequencing and Genomic Technologies research core facility. We trimmed and quality-filtered raw reads using *fastp* version 0.23.2 (Chen et al., 2018), mapped quality-filtered reads to the *P. eatonii* reference genome (Jarvis et al., 2025) using *bwa mem* version 0.7.17 (Li, 2013), and called genotypes to generate a filtered all-sites vcf file using *bcftools* version 1.18 (Li, 2011). See supplemental methods for additional details.

Raw sequencing data has been deposited in NCBI Sequence Read Archive (PRJNA1479764).

### Tree inference

We generated two sets of sequence alignments: for 10 kb non-overlapping genomic windows and for coding sequences (CDS), using custom python scripts reported in Stone and Wessinger (2024). We inferred a tree for each sequence alignment using *IQ-TREE* version 2.1.2 (Nguyen et al., 2015) and used *ASTRAL-III* version 5.7.1 (Zhang et al., 2018) to infer a species tree from the collection of 10 kb window or CDS region trees. We rooted each tree using the outgroup taxon, *P. palmeri*. We calculated gene concordance factors in *IQ-TREE* on each species tree (Minh et al., 2020). We inferred relative divergence times for the 10 kb window-based species tree using a penalized likelihood approach implemented in *treePL* version 1.0 (Smith & O’Meara, 2012). The resulting ultrametric tree was used for all phylogenetic comparative analyses. See supplemental methods for additional details on phylogenetic inference.

### Phylogenetic comparative analyses

To visualize variation in multi-trait phenotypes, we produced phylomorphospace plots (Sidlauskas, 2008) using the *phylomorphospace* function in the R package *phytools* (Revell, 2012). We used phylogenetic ANOVAs to quantify associations between each measured floral trait and pollination mode, accounting for shared phylogenetic history, using the *pgls* command of the R package *caper* (Orme et al., 2013) and specifying the correlation structure based on Pagel’s lambda (Pagel, 1994) to be estimated using maximum likelihood. We also investigated whether bee-adapted species that are sister to hummingbird-adapted species differed consistently in any measured floral trait compared to bee-adapted species that are not sister to hummingbird-adapted species. We scored all bee syndrome species as either sister or non-sister to hummingbird syndrome species and performed a phylogenetic ANOVA to test for differences in measured traits. We also identified evolutionary associations between pairs of measured traits across bee syndrome accessions using phylogenetic generalized least squares implemented using the *pgls* function of *caper*.

### Tests of phenotypic convergence

We quantified convergence among hummingbird syndrome taxa using the C1 convergence metric (Stayton, 2015). This metric is the ratio of the phenotypic distance between focal taxa relative to the maximum phenotypic distance since their evolutionary divergence, inferred using ancestral state reconstruction under a Brownian Motion model. Here, our focal taxa were the hummingbird syndrome tips. We performed this calculation using the *convrat* function of the R package *convevol* (Brightly & Stayton, 2023). To assess significance, we compared the calculated C1 value to 1000 evolutionary simulations using the *convratsig* function of *convevol*.

### Within-species variation in floral dimensions

Using digital calipers, we measured floral tube length, tube width, and stamen filament length of flowers sampled from natural populations. We collected measurements on 2 flowers per individual for up to 10 individuals per population for at least four populations per species for *P. barbatus*, *P. eatonii*, *P. laevis*, and *P. virgatus*. Population and locations measured can be found in Table S2. For each species, we tested for significant associations among average phenotype values by fitting linear models using the R package *nlme* (Pinheiro et al., 2017), with population included as a random effect.

### Clade-wide patterns of introgression

We used a framework based on *D*-statistics to examine clade-wide patterns of allele sharing between species due to introgression, using SNP data and the 10 kb-window *ASTRAL-III* tree as our species tree hypothesis. *D*-statistics quantify ratios of discordant topologies to infer introgression events using site patterns in sequence data (reviewed by Hibbins & Hahn, 2022). We filtered our vcf file for biallelic sites with a minor allele count >3 using *vcftools*. We calculated *D*-statistics and F4 ratios for all four-taxon subtrees using the *Dtrios* command in *Dsuite* version 0.5 (Malinsky et al., 2021). We then then used the *fbranch* command in *Dsuite* to assign introgression events to specific internal branches on the species tree. We adjusted the resulting *p*-values of the *f-branch* tests for multiple testing using the “Holm” method with alpha=0.001, implemented using the *p.adjust* function in R.

### Tests of introgression between pairs of hummingbird syndrome species

Each of the four hummingbird syndrome species has a bee syndrome sister taxon, a feature well-suited to tests of introgression using a 5-taxon rooted quartets framework: ((B1, H1), (B2, H2), O), where “B” denotes bee syndrome, “H” denotes hummingbird syndrome, “O” denotes the outgroup (*P. palmeri*), and numbers index the sister species pair. We used the *D_FOIL_* statistic (Pease & Hahn, 2015) to test for introgression between all six possible pairs of hummingbird syndrome taxa. For this analysis, we chose a single representative accession for each taxon based on highest sequence coverage (see Table S1 for a list of individuals used in this analysis), which is sufficient sampling to detect introgression between species (Hibbins & Hahn, 2022). For the genetically complicated *P. barbatus–P. virgatus* species complex, we chose a *P. barbatus* individual from the eastern clade and a *P. virgatus* individual from the western clade. We implemented the *D_FOIL_* tests using 100 kb window alignments, following the recommended best practices (Pease & Hahn, 2015).

For each of the six rooted quartets, we compared coalescence times in genomic windows showing the discordant syndrome topology (hummingbird syndrome taxa are sister) to coalescence times in windows showing the concordant species tree topology. First, we analyzed topological variation across the genome for each rooted quartet using the 10 kb window tree set. 15 possible topologies are possible for a given rooted quartet. Three of these support (B1, H1) as sister taxa, three topologies support (B2, H2) as sister taxa, and three support the discordant pattern where (H1, H2) are sister taxa. We used the program *Twisst* version 0.2 (Martin & Van Belleghem, 2017) to assign each 10 kb window tree to a given topology. As a proxy for coalescence time between focal individuals, we calculated relative node depth (RND), a normalized pairwise genetic distance (*d_XY_*) between focal taxa that controls for variation in mutation rate by normalizing to divergence from an outgroup. We first estimated pairwise genetic distance (*d_XY_*) between focal taxa in our windows using *pixy* version 1.2.7.beta1 (Korunes & Samuk, 2021). We then calculated relative node depth (RND) by dividing *d_XY_* between focal taxa by the average *d_XY_* between each taxon and the outgroup, *P. palmeri*. We then compared distributions of RND values across topologies using Mann-Whitney U tests (Suvorov et al., 2022).

### Effects of the choice of reference genome on results

Our primary analyses employ the *P. eatonii* reference genome (Jarvis et al., 2025), which is newer and higher quality than the other available reference genome in the section (for *P. barbatus*; Wessinger et al., 2023). To determine whether the choice of reference genome impacts the results of our genomic analyses, we repeated all steps using the *P. barbatus* genome for comparison.

## Results

### Hummingbird syndrome has evolved four times within *Penstemon* sect. *Habroanthus*

The inferred species trees based on 10 kb genomic windows and the CDS gene trees were nearly identical, differing only in the phylogenetic placements of *P. leiophyllus* and *P. wardii* (Figs. 1, S1). Despite the consistency between species trees, our gene concordance factor analysis revealed that most nodes in our tree are supported by just a small fraction of gene trees (Fig. 1). In cases where we sampled multiple accessions per species, they were nearly always monophyletic in the species tree. The sole exception to this pattern was the genetically intermingled *P. barbatus–P. virgatus* species complex. Consistent with prior work in this species complex (Wessinger et al., 2023), we identified two clades within each species that correspond to geographic origin: western accessions from Arizona vs. eastern accessions from Colorado and New Mexico. The eastern *P. barbatus* clade is more closely related to the eastern *P. virgatus* clade than it is to the western *P. barbatus* clade, reflecting interspecific gene flow within each geographic region.

**Figure 1.**
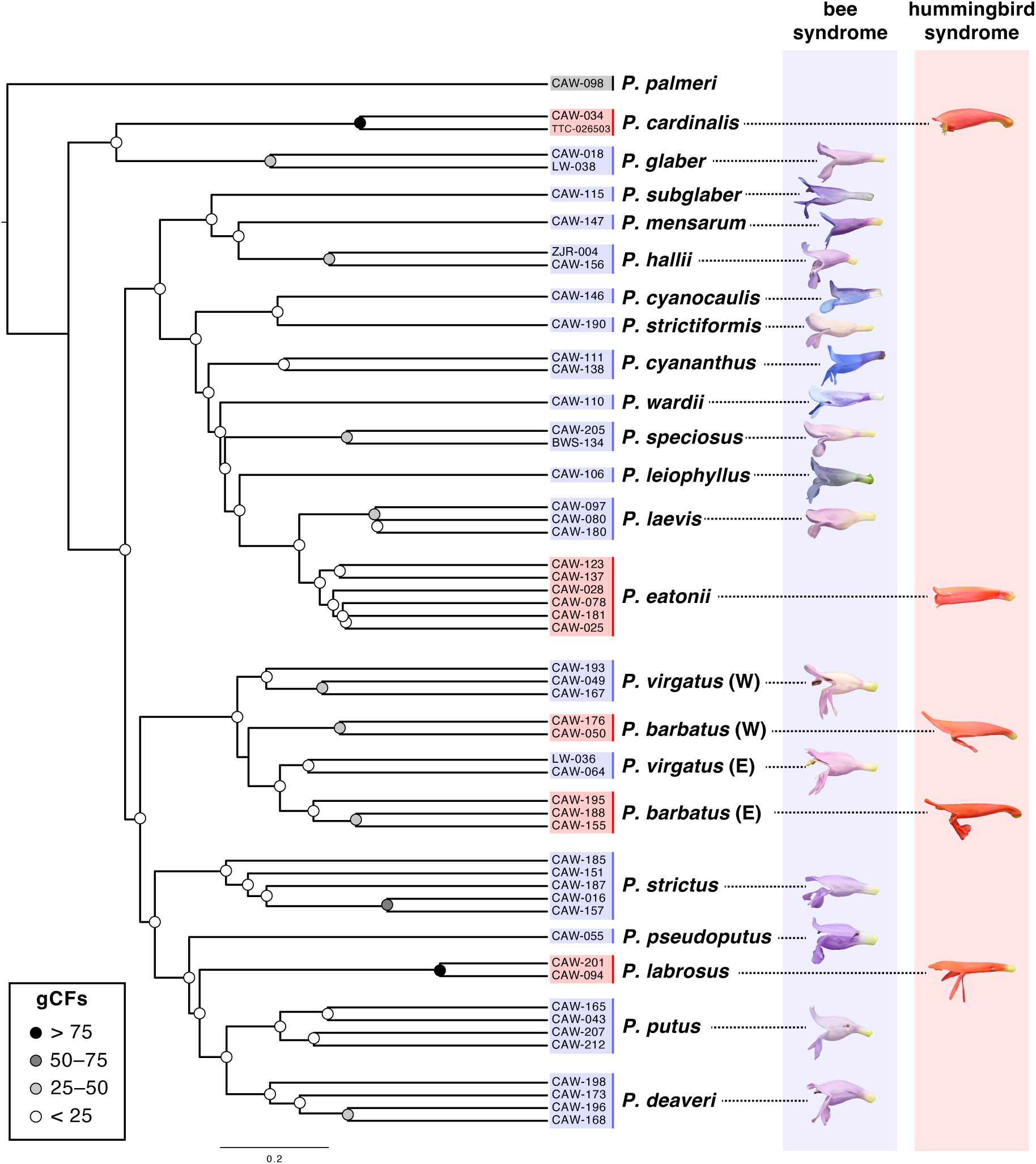
Phylogenetic relationships between sampled species in *Penstemon* section *Habroanthus* were inferred using a species tree approach based on 10 kb genomic windows. Node labels indicate gene concordance factors (gCFs), indicating the fraction of decisive gene trees supporting each branch. Paraphyletic lineages of *P. barbatus* and *P. virgatus* are labelled as western (W) or eastern (E).

Our phylogenetic estimates confirm that the four hummingbird-adapted species represent separate origins of this syndrome, with each of these species sister to a bee-adapted lineage (Fig. 1). These are (1) *P. barbatus–P. virgatus*, (2) *P. cardinalis–P. glaber*, (3) *P. eatonii–P. laevis*, and (4) *P. labrosus–P. deaveri* + *P. putus.* These species relationships allow us to take a sister species pair approach in our comparative methods and phylogenomic analyses.

### Convergent floral trait shifts accompany the evolution of hummingbird pollination

To test whether sampled accessions separate into distinct clusters in multivariate trait space that resemble the classic syndromes, we visualized patterns of floral trait variation across accessions using a PCA based on floral dimensions, nectar volume, nectary area, and anthocyanin type (Fig. 2A). The major axis of variation (PC1) explained 81% of the variation in traits across samples. This axis separated bee syndrome species (with delphinidin-based anthocyanins, small nectar volume, small nectaries, short floral tubes, and short reproductive organs) from hummingbird syndrome species (with pelargonidin- or cyanidin-based anthocyanins, large nectar volume, large nectaries, long and narrow floral tubes, and long reproductive organs). The second most important axis of variation (PC2) explained 9% of the variation and separated accessions with larger overall floral dimensions and more nectar from those with smaller flowers and less nectar.

**Figure 2.**
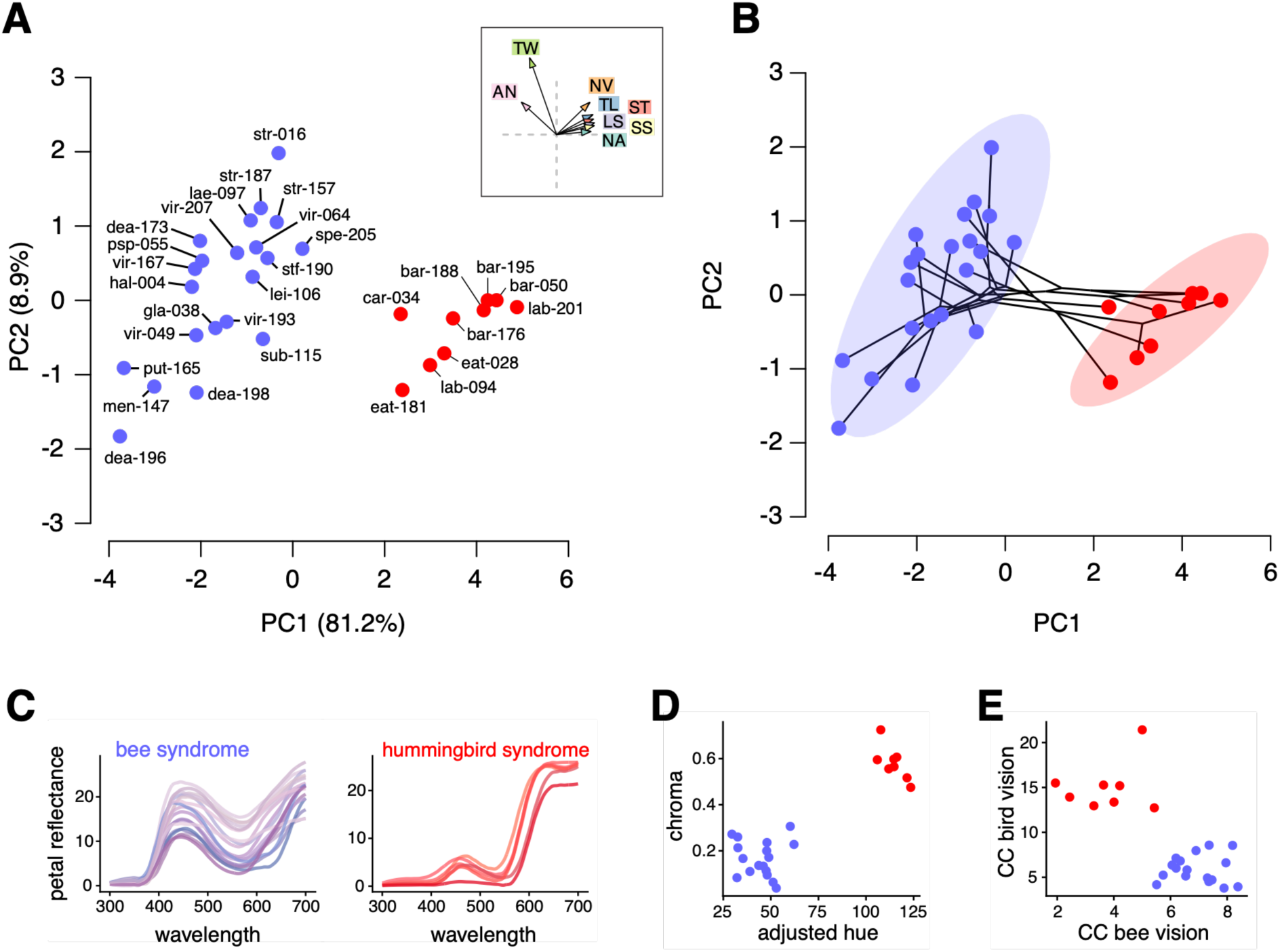
Multi-trait phenotypic differences between bee and hummingbird syndrome *Habroanthus* species. (A) Principal components analysis based on multiple traits, with accessions labeled according to population number as in Figure 1. Inset shows trait loadings onto PC axes. (B) Phylomorphospace plot based on the multi-trait PCA. (C) Petal reflectance spectra for bee and hummingbird syndrome flowers. (D) Relationship between adjusted hue and chroma values, calculated from petal reflectance data. (E) Relationship between chromatic contrast under the honeybee visual model (CC bee vision) and chromatic contrast under the avian violet sensitive model (CC bird vision), calculated from petal reflectance data. Purple denotes bee syndrome taxa and red denotes hummingbird syndrome taxa in all panels. AN: qualitative anthocyanin content, CC: chromatic contrast, LS: long stamen length, NA: log nectary area, NV: log nectar volume, SS: short stamen length, ST: style length, TL: floral tube length, TW: tube width. Abbreviations for accessions shown in panel A can be found in Table S1.

Our phylomorphospace plot confirms that sampled *Habroanthus* accessions separate into two pollination syndromes, and that the four hummingbird syndrome species have independently assembled trait combinations that map to a similar, circumscribed region of multi-trait space (Fig. 2B). We identified significant morphological convergence using the C1 statistic (C1 = 0.512, *p* < 0.001), suggesting hummingbird syndrome lineages have evolved to be more similar to each other than were their ancestors – they have shortened about half of the ancestral phenotypic distance through convergent evolution.

Bee syndrome species in *Penstemon* section *Habroanthus* have bluish-purple to pale flowers that are colored by blue delphinidin-based anthocyanins, sometimes in combination with pinkish cyanidin-based anthocyanins (Fig. 2C). Evolutionary shifts to red hummingbird syndrome flowers involve a shift to producing red pelargonidin-based pigments (*P. barbatus*, *P. eatonii*, and *P. labrosus*) or producing cyanidin pigments (*P. cardinalis*). The red flowers of all four hummingbird syndrome species exhibit broadly similar reflectance spectra (Fig. 2C), suggesting either cyanidin or pelargonidin can create similarly red colors, potentially in combination with yellow carotenoid pigments that are common secondary pigments in *Penstemon* flowers (León-Osper et al., 2025). In addition to shifting towards redder hue values, the four hummingbird syndrome species have convergently evolved increased petal chroma, in other words a more intensely saturated color (Fig. 2D; Table S3). The convergent flower color shifts to these chromatically intense red flowers yields high chromatic contrast under an avian visual model and reduced chromatic contrast under a honeybee visual model (Fig. 2E; Table S3).

### Floral syndrome traits exhibit lability and assemble rapidly during shifts in floral syndrome

An open question is whether shifts in specific floral traits, for example increased nectar reward or attractive colors, precede the evolution of hummingbird adaptation. In this scenario, bee-adapted species sister to hummingbird-adapted species will have evolved one or more initial steps towards hummingbird syndrome compared to other non-sister bee syndrome species. We tested this prediction by comparing trait values between these categories of species using a phylogenetic ANOVA. All measured traits differed significantly between bee and hummingbird syndrome taxa (Fig. 3A; Table S3). Because hummingbird syndrome flowers were both longer and narrower than bee syndrome flowers, we analyzed the ratio of floral tube length over width and found this ratio has a stronger association with syndrome than found for either trait individually (Fig. 3A; Table S3). Trait values for bee syndrome taxa did not differ according to their “sister status” – those taxa sister to hummingbird-adapted taxa did not have trait values more similar to the bird-adapted taxa compared to other bee-adapted taxa (Fig. 3A; Table S4).

**Figure 3.**
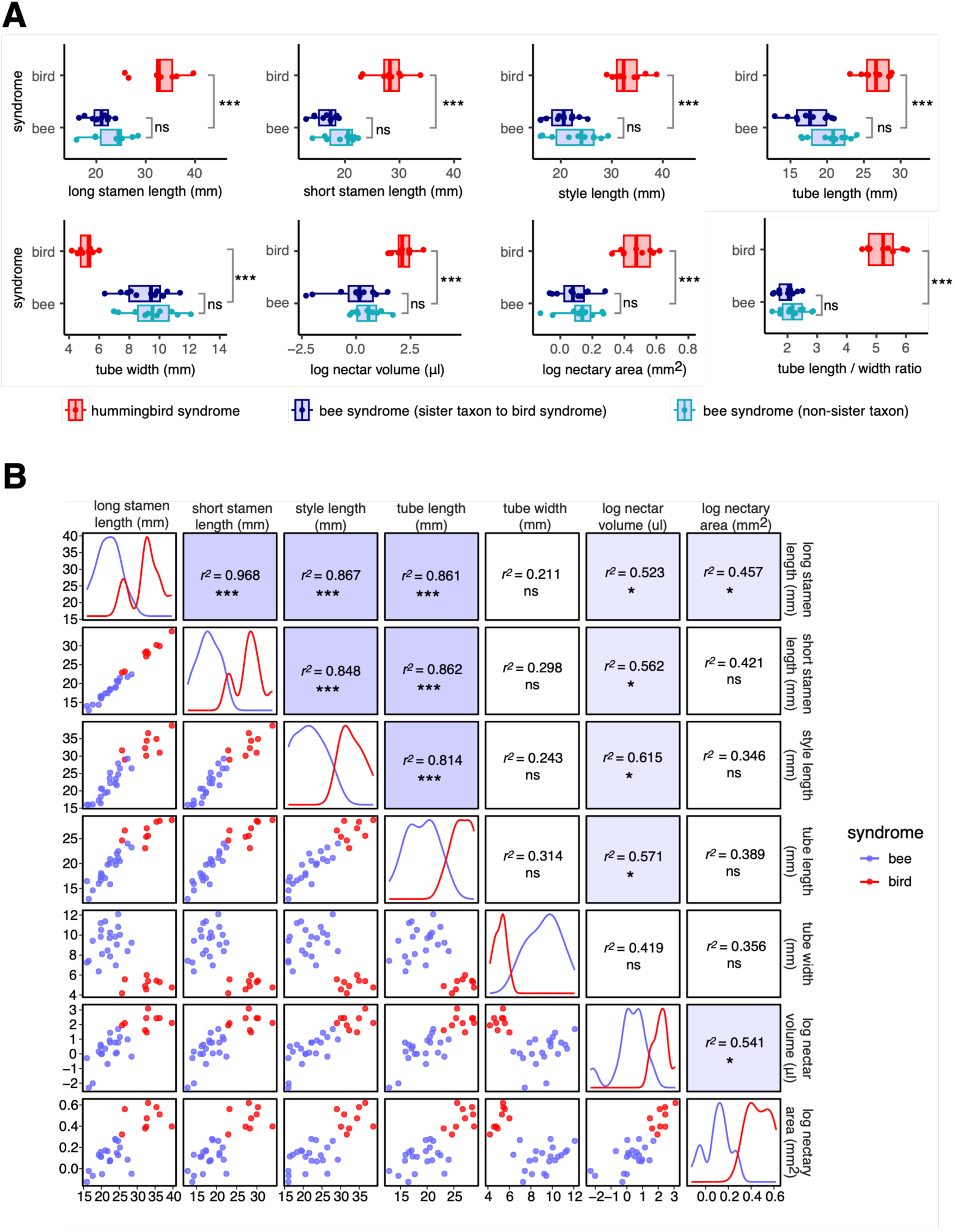
Floral trait associations within and between pollination syndromes. (A) Mean floral trait values for accessions across categories, showing results of phylogenetic ANOVAs testing for significant differences between bee vs. hummingbird syndrome and between sister vs. non-sister bee syndrome species. (B) Pairwise correlations among floral traits with phylogenetic correlations among bee syndrome species reported. *: *p* < 0.05, ***: *p* < 0.0001, ns: *p* > 0.05.

To examine whether patterns of floral trait variation in bee syndrome taxa might facilitate or constrain shifts to hummingbird syndrome, we estimated evolutionary correlations among floral traits across the set of bee syndrome taxa using phylogenetic generalized linear regression models. Among bee syndrome taxa, floral organ lengths (petal tube, stamens, and style) showed very strong positive evolutionary correlations (Fig 3B; Table S5). Floral tube width was not correlated with any measured floral trait, suggesting this trait has evolved independently of overall flower size. Nectar volume was only weakly correlated with floral length traits and nectary area. For four species (the bee syndrome *P. laevis* and *P. virgatus* and the hummingbird syndrome *P. barbatus* and *P. eatonii*), we measured floral dimension phenotypes in natural populations to estimate phenotypic correlations. All species exhibited strong linear relationships between floral organ length traits and moderate or no relationship between organ length and floral tube width (Fig S2).

### *Penstemon* sect. *Habroanthus* has a history of introgression and incomplete lineage sorting

The relatively low gene concordance factors in our species tree analysis suggests significant variation in the history of coalescence across the genome. Even the deepest nodes in our tree showed substantial discordance (Fig. 1). Both incomplete lineage sorting (ILS) and introgression following hybridization events can generate such discordance. Our *f*-branch analysis characterized patterns of introgression across the species tree, finding evidence that introgression has been prevalent in the evolutionary history of section *Habroanthus*, particularly between pairs of bee syndrome species collected in geographic proximity (Fig S3). Examples of these cases include *P. glaber* and *P. hallii* collected in Colorado; *P. cyananthus*, *P. subglaber*, and *P. wardii* collected in Utah; and *P. leiophyllus* with *P. strictus*, *P. pseudoputus*, *P. putus*, and *P. deaveri*, all collected in Utah and Arizona. In some cases, introgression events mapped to internal branches of the tree.

### No evidence for introgression fueling convergent origins of hummingbird syndrome

We tested for introgression between all six possible pairs of hummingbird syndrome taxa with rooted 5-taxon trees using the *D_FOIL_* framework. None of the four component *D*-statistics that form the *D_FOIL_* test were significantly different from zero in any of the six tests (Fig. 4A), indicating no genome-wide significant pattern of introgression is found between any of pair of hummingbird syndrome species.

**Figure 4.**
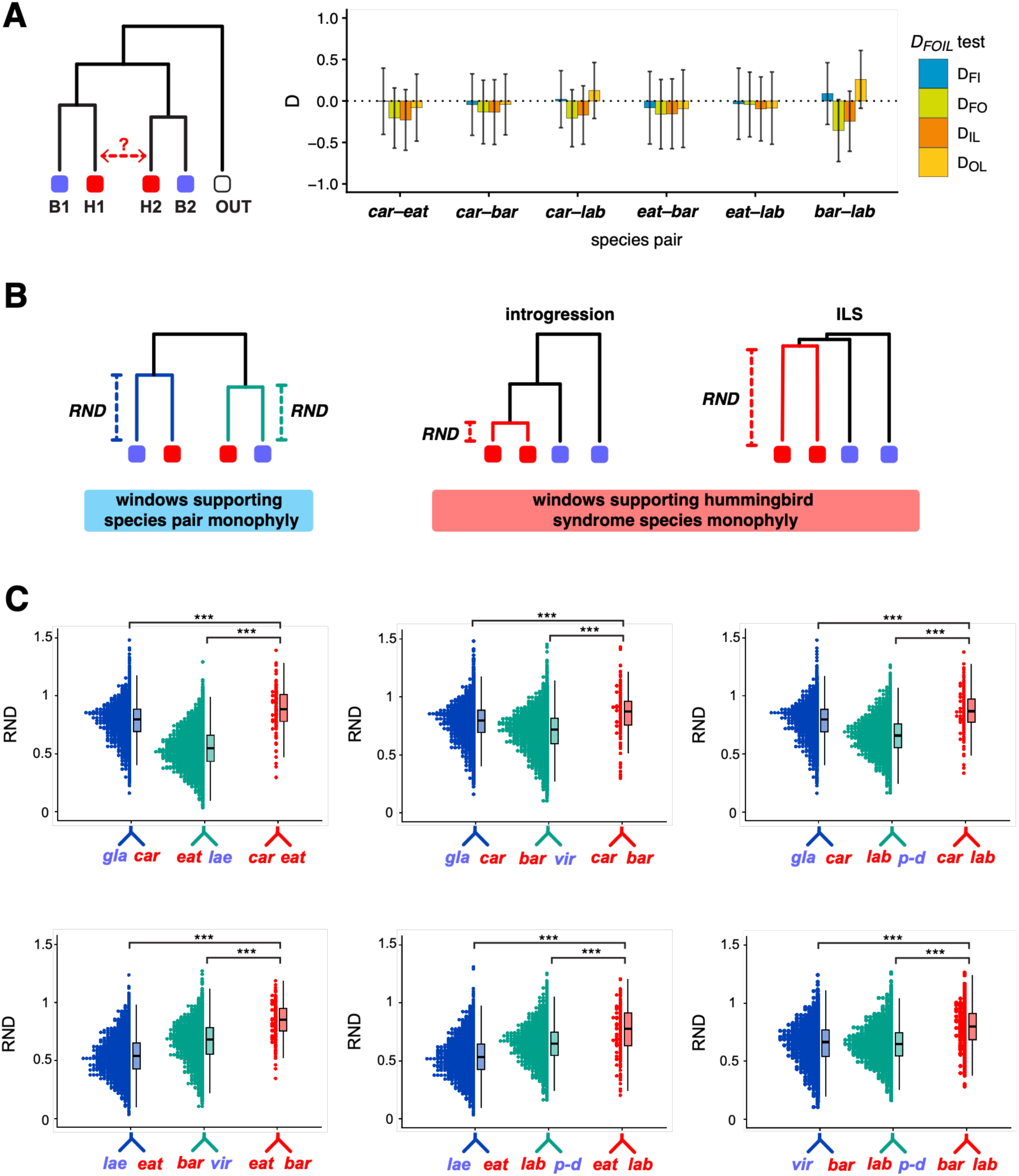
Results of 5-taxon tests for introgression. (A) Example 5-taxon rooted quartet and results of *D_FOIL_* tests for all six pairwise tests for introgression. (B) Conceptual logic of relative coalescence test for introgression including predicted relative node depth (RND) values under hypotheses of introgression vs. incomplete lineage sorting (ILS). (C) Distributions of relative node depth (RND) values between sister species for genomic windows concordant with the species tree topology for species pair 1 and 2 (navy and green, respectively) and for pairs of hummingbird syndrome taxa in genomic regions supporting allele sharing between them (red). Significance of Mann-Whitney U tests are reported. bar: *P. barbatus*, car: *P. cardinalis*, eat: *P. eatonii*, gla: *P. glaber*, lab: *P. labrosus*, lae: *P. laevis*, p-d: *P. putus + P. deaveri*, and vir: *P. virgatus*. ***: *p* < 0.0001.

Despite this lack of genome-wide significance, we examined patterns of sequence divergence in regions of the genome showing allele sharing between each pair of hummingbird syndrome species. In theory, these regions could arise either due to a history of introgression between hummingbird syndrome species that failed to leave a genome-wide signal or due to ILS. These alternatives can be distinguished based on relative sequence coalescence times. Introgression should generate genomic regions with recent coalescence between hummingbird syndrome sequences – more recent than expected under the species tree concordant topology with complete lineage sorting (Fig. 4B). Alternatively, allele sharing between hummingbird syndrome species due to ILS should result in coalescence at least as deep as that expected under the concordant topology (Fig. 4B). For each of the six species pair comparisons, we found significantly deeper coalescence times (large RND values) between hummingbird syndrome species in windows showing allele sharing, compared to coalescence times between hummingbird syndrome species and their sister bee syndrome taxon in the species tree concordant windows (Fig. 4C; Table S6). This deeper coalescence time matches observed values in genomic regions supporting other discordant topologies (Fig. S4; Table S6), and overall is consistent with ILS causing discordant genealogies.

Our choice of reference genome has no impact on our results – when we repeated all phylogenetic analyses using the *P. barbatus* reference, we found identical conclusions (Figs. S5-S8; Tables S7-S10).

## Discussion

Our updated phylogeny for *Penstemon* section *Habroanthus* reveals an evolutionary history consistent with a burst of diversification, rapid floral evolution, and incomplete sorting of ancestral variation. We found that floral syndrome is approximately binary in this section, with no clearly intermediate taxa. The four hummingbird-adapted taxa represent four separate origins of this syndrome (Fig. 1), making this clade a useful case study for understanding how and why hummingbird pollination has evolved so many times across the genus and other western North American perennial wildflower lineages (Abrahamczyk & Renner, 2015; Grant, 1994; Stebbins, 1989; Wilson et al., 2007). Species with bee-adapted flowers show floral morphological diversity, consistent with adaptation on distinct assemblages of bee and wasp pollinators (Wilson et al., 2004). The four hummingbird syndrome species exhibit significant convergence in their multi-trait phenotypes (Fig. 2B), suggesting that lineages are drawn towards similar adaptive optima from distinct ancestral phenotypes during adaptation to hummingbirds. These evolutionary transitions occurred relatively recently, suggesting evolutionary lability towards hummingbird syndrome in this adaptive radiation (Thomson & Wilson, 2008).

Red or reddish flower color is the most reliable indicator of hummingbird adaptation in penstemons (Wilson et al., 2004). We found that convergence on a similar red color, with high chromatic contrast to bird visual systems and low chromatic contrast to bee visual systems, has occurred through different types of anthocyanin pigments in section *Habroanthus* (Fig. 2C-E). The evolution of red flowers through distinct pigment combinations has also been documented in the family Solanaceae (Ng & Smith, 2016) and in a comparative dataset across various hummingbird-adapted Californian taxa (León-Osper et al., 2025). Convergence on red flowers by hummingbird-adapted plants in western North America likely reflects selection for mimicry of a common signal highly detectable by migratory hummingbirds (Brown & Kodric-Brown, 1979; Crosswhite & Crosswhite, 1981; Grant, 1966) and perhaps also selection to reduce detectability to bees (Chittka et al., 2001). Thus, the evolution of red colors that both maximize detectability to birds and minimize detectability of bees suggests both pro-bird and anti-bee adaptation (de Camargo et al., 2019; León-Osper et al., 2025; Ng & Smith, 2016; Ogutcen et al., 2020). However, we note that pollinator behavior cannot be directly extrapolated from models of chromatic contrast. For example, prior observations of pollinator visitation suggest substantial bee visitation to hummingbird syndrome *Penstemon* species (e.g., Kimball, 2008; Wilson et al., 2004).

### No evidence for gradual, stepwise assembly of hummingbird syndrome

A goal of this study was to test whether ancestral phenotypes in *Penstemon* jumpstart shifts to hummingbird syndrome, by reducing the number of subsequent changes. There may be an order to trait evolution, where the evolution of increased hummingbird attraction should initiate a transition to hummingbird syndrome, followed by the evolution of morphological traits that fine-tune the efficiency of pollen transfer (Armbruster, 2014; Stebbins, 1970; Stebbins, 1974). In particular, the evolution of generous nectar offerings has been considered to be a prerequisite to evolution of hummingbird syndrome flowers in *Penstemon* (Thomson & Wilson, 2008; Wilson et al., 2006). Indeed, a manipulative ecological study found that hummingbirds increase their visitation rate to bee-adapted *Penstemon* flowers that are supplemented with greater nectar volume (Wilson & Jordan, 2009).

However, we found no evidence that the evolution of increased nectar – or any other hummingbird syndrome trait – preceded the most recent speciation events. Bee syndrome taxa sister to hummingbird syndrome taxa tended to have similar trait values to other bee syndrome species (Fig. 3A). Instead, it appears that the hummingbird syndrome is assembled rapidly on the lineage leading to each hummingbird syndrome species. This may have involved an ordered series of steps, however we are unable to discern any order to trait evolution and cannot evaluate the hypothesis that shifts in reward or attraction traits precede the evolution of morphological traits. We note that it remains possible that the ancestor to both species had large nectar volume that was subsequently reduced in the bee syndrome sister species, or that bee syndrome sister taxa harbor population-level variation in nectar production that was not captured by our sampling.

### Patterns of correlations among traits across and within bee-adapted species may facilitate shifts to hummingbird syndrome

Developmental and genetic constraints on the production of phenotypic variation can shape the evolution of multi-trait phenotypes (Lande & Arnold, 1983; Maynard Smith et al., 1985; Schluter, 1996). To explore potential biases on the production of variation, we examined patterns of correlations among floral traits in bee syndrome species – the ancestral character state. In agreement with a previous morphological analysis of a largely different set of *Penstemon* species than sampled here (Wilson et al., 2004), we found that bee syndrome species vary substantially in their multi-trait floral phenotypes. Across bee syndrome species, we identified strong positive evolutionary correlations between floral organ length traits (corolla, stamen filaments, and styles) and weak to no correlation between organ length traits and tube width or between nectar traits and floral dimensions (Fig. 3B). Overall, these findings suggest that bee syndrome species exhibit diverse combinations of phenotypes with few obvious constraints on the production of variation that is relevant to shifts to hummingbird pollination.

The evolutionary correlations among floral dimensions in bee syndrome species resemble patterns of trait variation observed within natural populations. Our population-level measurements include the effects of environmental variation on trait values, and so they are not estimates of genetic correlations. However, we find striking evidence of a much stronger positive correlation between floral organ length traits than between floral tube length and width in both bee- and hummingbird-adapted populations (Fig. S2). These observations match patterns of phenotypic variation measured in a larger population of *P. virgatus* (Wessinger et al., 2018) and in natural populations of five other bee-adapted penstemons (Rodríguez-Peña & Wolfe, 2023). The positive correlation between organ length traits is parallel to the direction of selection during an evolutionary shift to hummingbird syndrome – a feature that may facilitate shifts between syndromes (Wessinger & Hileman, 2016). On the other hand, largely decoupled floral length and width traits suggests that relatively long and narrow floral tubes with exserted reproductive organs of hummingbird syndrome species are evolutionarily accessible in *Penstemon*.

### No evidence for introgression fueling repeated shifts to hummingbird syndrome

Adaptive introgression is a compelling mechanism for rapid shifts in complex traits during species radiations. In this scenario, a hybridization event, followed by repeated backcrossing to parental species, transfers adaptive alleles across species boundaries. Hybridization can be common in radiations if reproductive barriers are incomplete and lineages are geographically proximal. In fact, many iconic radiations are characterized by introgression, which has allowed lineages to repeatedly access novel ecological niches (Baiz et al., 2021; Edelman et al., 2019; Lamichhaney et al., 2015; Meier et al., 2017; Richards & Martin, 2017; Wogan et al., 2023). Stebbins (1989), noting the lack of strong reproductive isolating barriers and prevalence of hybridization in *Penstemon*, speculated that hummingbird syndrome alleles have moved between lineages through adaptive introgression.

Our phylogenomic analyses uncovered pervasive gene tree discordance due to ILS, matching every prior phylogenomic study of *Penstemon* to date (e.g., Depatie & Wessinger, 2025; Stone & Wessinger, 2024; Stone & Wolfe, 2021; Wessinger et al., 2016; Wolfe et al., 2006). We also identified evidence of introgression between geographically proximal bee syndrome species (Fig. S3). However, we found no evidence that introgression has fueled repeated origins to hummingbird syndrome in *Penstemon*. Allele sharing between hummingbird syndrome taxa was best explained by ILS, not introgression (Fig. 4).

While we found no evidence for adaptive introgression, it remains possible that selection has repeatedly targeted the same genes through de novo mutations. This is the case for flower color, where the evolution of red from blue-purple involves de novo loss-of-function mutations to the anthocyanin pathway gene *Flavonoid 3’,5’-hydroxylase*, which normally functions to produce blue delphinidin-based pigments (Stone & Wessinger, 2024; Wessinger & Rausher, 2015). Adaptation from standing genetic variation may also contribute to repeated evolution and has recently been identified in diverse study systems (e.g., Fuhrmann et al., 2023; Meier et al., 2018; Roberts Kingman et al., 2021; Rubin et al., 2022). At present, we cannot evaluate the contribution of gene re-use or adaptation from ancestral variation because we currently lack datasets identifying the genomic loci for hummingbird pollination across all four origins.

## Conclusions

To understand evolved features that predispose *Penstemon* towards shifts to hummingbird syndrome, we may need to take a broader view of traits shared by the genus. *Penstemon* is a perennial genus, a life history strategy shared by (nearly) all hummingbird-adapted plants in western North America flora, which tend to flower over long time periods in the summer months (Grant & Grant, 1968; Stebbins, 1989). All penstemons have flowers with a bilateral tubular shape that can be modified into the narrowly tubular flowers adapted for hummingbird pollination (Castellanos et al., 2004). The nectaries of *Penstemon* flowers replenish nectar throughout the day, helping them entrain hummingbird pollinators to regular visits (Thomson & Wilson, 2008). Most species lack a strong floral scent, making transition to moth pollination seemingly difficult. Simple loss-of-function mutations to anthocyanin pathway genes result in evolutionary shifts from blue-purple to red flowers across the genus (Wessinger & Rausher, 2015). Finally, a capacity for self-fertilization may also facilitate specialization on hummingbirds as pollinators (Wessinger & Kelly, 2018). Transitions to hummingbird pollination surely begin with a shift in environmental conditions favored by hummingbirds (Stebbins, 1989; Thomson & Wilson, 2008). For western North American penstemons, these habitats tend to be at lower elevations and lower latitudes (Hamilton & Wessinger, 2022). We conclude that the strong convergence between hummingbird syndrome species results from closely-related species responding in similar ways to similar environmental pressures (Blount et al., 2018).

## Data and code availability

Data and scripts to reproduce analyses will be posted on Zenodo following publication.

## Author contributions

Molecular work and phenotypic measurements were performed by NHW, PWL, and CAW. Data analyses were performed by CAW, BWS, LCW, and PWL. CAW funded this work and wrote the manuscript, with edits from all authors.

## Acknowledgements

We thank Zack Radford for collecting a *P. hallii* sample used in this study and Craig Freeman for confirming species identifications. We also thank Lena Hileman, Brian Hollis, and members of the Wessinger lab for helpful feedback on this work. We gratefully acknowledge the computational resources provided by the Hyperion high performance computing cluster at the University of South Carolina. We also acknowledge the technical assistance and resources provided by Research Computing at the University of South Carolina (RRID:SCR_027488).

## References

Abrahamczyk, S., & Renner, S. S. (2015). The temporal build-up of hummingbird/plant mutualisms in North America and temperate South America. BMC evolutionary biology, 15(1), 104–116.

Armbruster, W. S. (2014). Floral specialization and angiosperm diversity: phenotypic divergence, fitness trade-offs and realized pollination accuracy. AoB Plants, 6, plu003.

Baiz, M. D., Wood, A. W., Brelsford, A., Lovette, I. J., & Toews, D. P. (2021). Pigmentation genes show evidence of repeated divergence and multiple bouts of introgression in Setophaga warblers. Current Biology, 31(3), 643–649. e643.

Bell, M. A., & Foster, S. A. (1994). Introduction to the evolutionary biology of the threespine stickleback. The evolutionary biology of the threespine stickleback, 1–27.

Beuttell, K., & Losos, J. B. (1999). Ecological morphology of Caribbean anoles. Herpetological Monographs, 1–28.

Blount, Z. D., Lenski, R. E., & Losos, J. B. (2018). Contingency and determinism in evolution: Replaying life’s tape. Science, 362(6415), eaam5979.

Boccia, C. K., Swierk, L., Ayala-Varela, F. P., Boccia, J., Borges, I. L., Estupiñán, C. A., Martin, A. M., Martínez-Grimaldo, R. E., Ovalle, S., & Senthivasan, S. (2021). Repeated evolution of underwater rebreathing in diving Anolis lizards. Current Biology, 31(13), 2947–2954. e2944.

Brightly, W., & Stayton, C. (2023). convevol: Analysis of convergent evolution. R package version, 2(1).

Brower, A. V. (1996). Parallel race formation and the evolution of mimicry in Heliconius butterflies: a phylogenetic hypothesis from mitochondrial DNA sequences. Evolution, 50(1), 195–221.

Brown, J. H., & Kodric-Brown, A. (1979). Convergence, competition, and mimicry in a temperate community of hummingbird-pollinated flowers. Ecology, 60(5), 1022–1035.

Burgess, S. J., & Hibberd, J. M. (2015). Insights into C4 metabolism from comparative deep sequencing. Current Opinion in Plant Biology, 25, 138–144.

Castellanos, M., Wilson, P., & Thomson, J. (2004). ‘Anti-bee’and ‘pro-bird’changes during the evolution of hummingbird pollination in Penstemon flowers. Journal of evolutionary biology, 17(4), 876–885.

Castellanos, M. C., Wilson, P., & Thomson, J. D. (2003). Pollen transfer by hummingbirds and bumblebees, and the divergence of pollination modes in *Penstemon*. Evolution, 57(12), 2742–2752.

Chan, Y. F., Marks, M. E., Jones, F. C., Villarreal, G., Shapiro, M. D., Brady, S. D., Southwick, A. M., Absher, D. M., Grimwood, J., & Schmutz, J. (2010). Adaptive evolution of pelvic reduction in sticklebacks by recurrent deletion of a Pitx1 enhancer. Science, 327(5963), 302–305.

Chen, S., Zhou, Y., Chen, Y., & Gu, J. (2018). fastp: an ultra-fast all-in-one FASTQ preprocessor. Bioinformatics, 34(17), i884–i890.

Chittka, L., Spaethe, J., Schmidt, A., & Hickelsberger, A. (2001). Adaptation, constraint, and chance in the evolution of flower color and pollinator color vision. In Cognitive ecology of pollination: animal behavior and floral evolution. Cambridge University Press.

Christin, P.-A., Osborne, C. P., Chatelet, D. S., Columbus, J. T., Besnard, G., Hodkinson, T. R., Garrison, L. M., Vorontsova, M. S., & Edwards, E. J. (2013). Anatomical enablers and the evolution of C4 photosynthesis in grasses. Proceedings of the National Academy of Sciences, 110(4), 1381–1386.

Colosimo, P. F., Hosemann, K. E., Balabhadra, S., Villarreal Jr, G., Dickson, M., Grimwood, J., Schmutz, J., Myers, R. M., Schluter, D., & Kingsley, D. M. (2005). Widespread parallel evolution in sticklebacks by repeated fixation of ectodysplasin alleles. Science, 307(5717), 1928–1933.

Crosswhite, F., & Crosswhite, C. (1981). Hummingbirds as pollinators of flowers in the red-yellow segment of the color spectrum, with special reference to Penstemon and the” open habitat.”.

Crosswhite, F. S. (1965). Hybridization of Penstemon barbatus (Scrophulariaceae) of section Elmigera with species of a section Habroanthus. The Southwestern Naturalist, 234–237.

Crump, W. W., Stettler, J. M., Johnson, R. L., Anderson, C. D., Harrison, S., Meservey, L. M., & Stevens, M. R. (2020). Flower Color Variation in Jones’ Penstemon, Penstemon× jonesii Pennell (P. eatonii A. Gray× P. laevis Pennell)(Plantaginaceae). Western North American Naturalist, 80(2), 131–145.

de Camargo, M. G. G., Lunau, K., Batalha, M. A., Brings, S., de Brito, V. L. G., & Morellato, L. P. C. (2019). How flower colour signals allure bees and hummingbirds: a community-level test of the bee avoidance hypothesis. New Phytologist, 222(2), 1112–1122.

de los Santos-Gómez, S. M., & Ornelas, J. F. (2025). Shared history between two Penstemon (Plantaginaceae) species: potential hybridization between populations: de los Santos-Gómez, Ornelas. Plant Systematics and Evolution, 311(5), 37.

Depatie, T. H., & Wessinger, C. A. (2025). The unique morphological basis and repeated evolutionary origins of personate flowers in Penstemon. American Journal of Botany, 112(8), e70078.

Dowling, T. E., Martasian, D. P., & Jeffery, W. R. (2002). Evidence for multiple genetic forms with similar eyeless phenotypes in the blind cavefish, Astyanax mexicanus. Molecular biology and evolution, 19(4), 446–455.

Edelman, N. B., Frandsen, P. B., Miyagi, M., Clavijo, B., Davey, J., Dikow, R. B., García-Accinelli, G., Van Belleghem, S. M., Patterson, N., & Neafsey, D. E. (2019). Genomic architecture and introgression shape a butterfly radiation. Science, 366(6465), 594–599.

Enbody, E. D., Sendell-Price, A. T., Sprehn, C. G., Rubin, C.-J., Visscher, P. M., Grant, B. R., Grant, P. R., & Andersson, L. (2023). Community-wide genome sequencing reveals 30 years of Darwin’s finch evolution. Science, 381(6665), eadf6218.

Freeman, C. C. (2019). Penstemon. In F. o. N. A. E. Committee (Ed.), Flora of North America North of Mexico (Vol. 17: Magnoliophyta: Tetrachondraceae to Orobanchaeceae, pp. 82–255). Oxford University Press.

Fuhrmann, N., Prakash, C., & Kaiser, T. S. (2023). Polygenic adaptation from standing genetic variation allows rapid ecotype formation. elife, 12, e82824.

Galen, S. C., Natarajan, C., Moriyama, H., Weber, R. E., Fago, A., Benham, P. M., Chavez, A. N., Cheviron, Z. A., Storz, J. F., & Witt, C. C. (2015). Contribution of a mutational hot spot to hemoglobin adaptation in high-altitude Andean house wrens. Proceedings of the National Academy of Sciences, 112(45), 13958–13963.

Gould, S. J., & Vrba, E. S. (1982). Exaptation—a missing term in the science of form. Paleobiology, 8(1), 4–15.

Grant, K. A. (1966). A hypothesis concerning the prevalence of red coloration in California hummingbird flowers. The American Naturalist, 100(911), 85–97.

Grant, K. A., & Grant, V. (1968). Hummingbirds and their flowers. Columbia University Press.

Grant, V. (1994). Historical development of ornithophily in the western North American flora. Proceedings of the National Academy of Sciences, 91(22), 10407–10411.

Hamilton, A. M., & Wessinger, C. A. (2022). Adaptation to lower latitudes and lower elevations precedes the evolution of hummingbird pollination in western North American Penstemon. American Journal of Botany, 109(6), 1047–1055.

Hibbins, M. S., & Hahn, M. W. (2022). Phylogenomic approaches to detecting and characterizing introgression. Genetics, 220(2), iyab173.

Hooper, D. M., McDiarmid, C. S., Powers, M. J., Justyn, N. M., Kučka, M., Hart, N. S., Hill, G. E., Andolfatto, P., Chan, Y. F., & Griffith, S. C. (2024). Spread of yellow-bill-color alleles favored by selection in the long-tailed finch hybrid system. Current Biology, 34(23), 5444–5456. e5448.

Jarvis, D. E., Stevens, M. R., Carter, P., Lin, Y. F., Jaggi, K. E., Jijon, G., Kalt, T., Calixto, J., Standring, S., & Torres, K. (2025). Whole-genome assembly and annotation of the firecracker penstemon (Penstemon eatonii). Journal of Heredity, 116(3), 373–381.

Jones, M. R., Mills, L. S., Alves, P. C., Callahan, C. M., Alves, J. M., Lafferty, D. J., Jiggins, F. M., Jensen, J. D., Melo-Ferreira, J., & Good, J. M. (2018). Adaptive introgression underlies polymorphic seasonal camouflage in snowshoe hares. Science, 360(6395), 1355–1358.

Kimball, S. (2008). Links between floral morphology and floral visitors along an elevational gradient in a Penstemon hybrid zone. Oikos, 117(7), 1064–1074.

Korunes, K. L., & Samuk, K. (2021). pixy: Unbiased estimation of nucleotide diversity and divergence in the presence of missing data. Molecular ecology resources, 21(4), 1359–1368.

Lamichhaney, S., Berglund, J., Almén, M. S., Maqbool, K., Grabherr, M., Martinez-Barrio, A., Promerová, M., Rubin, C.-J., Wang, C., & Zamani, N. (2015). Evolution of Darwin’s finches and their beaks revealed by genome sequencing. Nature, 518(7539), 371–375.

Lande, R., & Arnold, S. J. (1983). The measurement of selection on correlated characters. Evolution, 1210–1226.

León-Osper, M., Rossi, V., Conrad, K., Mena, J. H., Meslow, E., Fuller, A., Narbona, E., & Whittall, J. B. (2025). California red hummingbird flowers: color convergence across four biochemical categories. Madroño, 72(2), 68–95.

Li, H. (2011). A statistical framework for SNP calling, mutation discovery, association mapping and population genetical parameter estimation from sequencing data. Bioinformatics, 27(21), 2987–2993.

Li, H. (2013). Aligning sequence reads, clone sequences and assembly contigs with BWA-MEM. arXiv preprint arXiv:1303.3997.

Losos, J. B. (2011). Convergence, adaptation, and constraint. Evolution, 65(7), 1827–1840.

Malinsky, M., Matschiner, M., & Svardal, H. (2021). Dsuite-Fast D-statistics and related admixture evidence from VCF files. Molecular ecology resources, 21(2), 584–595.

Martin, S. H., & Van Belleghem, S. M. (2017). Exploring evolutionary relationships across the genome using topology weighting. Genetics, 206(1), 429–438.

Maynard Smith, J., Burian, R., Kauffman, S., Alberch, P., Campbell, J., Goodwin, B., Lande, R., Raup, D., & Wolpert, L. (1985). Developmental constraints and evolution. Quarterly Review of Biology, 60(3), 265–287.

McGlothlin, J. W., Kobiela, M. E., Wright, H. V., Mahler, D. L., Kolbe, J. J., Losos, J. B., & Brodie III, E. D. (2018). Adaptive radiation along a deeply conserved genetic line of least resistance in Anolis lizards. Evolution letters, 2(4), 310–322.

Meier, J. I., Marques, D. A., Mwaiko, S., Wagner, C. E., Excoffier, L., & Seehausen, O. (2017). Ancient hybridization fuels rapid cichlid fish adaptive radiations. Nature communications, 8(1), 1–11.

Meier, J. I., Marques, D. A., Wagner, C. E., Excoffier, L., & Seehausen, O. (2018). Genomics of parallel ecological speciation in Lake Victoria cichlids. Molecular biology and evolution, 35(6), 1489–1506.

Minh, B. Q., Hahn, M. W., & Lanfear, R. (2020). New methods to calculate concordance factors for phylogenomic datasets. Molecular biology and evolution, 37(9), 2727–2733.

Ng, J., & Smith, S. D. (2016). How to make a red flower: the combinatorial effect of pigments. AoB Plants, 8.

Nguyen, L.-T., Schmidt, H. A., Von Haeseler, A., & Minh, B. Q. (2015). IQ-TREE: a fast and effective stochastic algorithm for estimating maximum-likelihood phylogenies. Molecular biology and evolution, 32(1), 268–274.

Ogutcen, E., Durand, K., Wolowski, M., Clavijo, L., Graham, C., Glauser, G., & Perret, M. (2020). Chemical basis of floral color signals in Gesneriaceae: the effect of alternative anthocyanin pathways. Frontiers in plant science, 11, 604389.

Orme, D., Freckleton, R., Thomas, G., Petzoldt, T., Fritz, S., Isaac, N., & Pearse, W. (2013). The caper package: comparative analysis of phylogenetics and evolution in R. R package version, 5(2), 1–36.

Pagel, M. (1994). Detecting correlated evolution on phylogenies: a general method for the comparative analysis of discrete characters. Proceedings of the Royal Society of London B: Biological Sciences, 255(1342), 37–45.

Pardo-Diaz, C., Salazar, C., Baxter, S. W., Merot, C., Figueiredo-Ready, W., Joron, M., McMillan, W. O., & Jiggins, C. D. (2012). Adaptive introgression across species boundaries in Heliconius butterflies. PLoS genetics, 8(6), e1002752.

Pease, J. B., & Hahn, M. W. (2015). Detection and polarization of introgression in a five-taxon phylogeny. Systematic biology, 64(4), 651–662.

Pinheiro, J., Bates, D., DebRoy, S., Sarkar, D., Heisterkamp, S., Van Willigen, B., & Maintainer, R. (2017). Package ‘nlme’. Linear and nonlinear mixed effects models, version, 3(1).

Revell, L. J. (2012). phytools: an R package for phylogenetic comparative biology (and other things). Methods in Ecology and Evolution, 3(2), 217–223.

Richards, E. J., & Martin, C. H. (2017). Adaptive introgression from distant Caribbean islands contributed to the diversification of a microendemic adaptive radiation of trophic specialist pupfishes. PLoS genetics, 13(8).

Roberts Kingman, G. A., Vyas, D. N., Jones, F. C., Brady, S. D., Chen, H. I., Reid, K., Milhaven, M., Bertino, T. S., Aguirre, W. E., & Heins, D. C. (2021). Predicting future from past: The genomic basis of recurrent and rapid stickleback evolution. Science Advances, 7(25), eabg5285.

Rodríguez-Peña, R. A., & Wolfe, A. D. (2023). Flower morphology variation in five species of Penstemon (Plantaginaceae) displaying Hymenoptera pollination syndrome. Botanical Sciences, 101(1), 217–232.

Rossi, M., Hausmann, A. E., Alcami, P., Moest, M., Roussou, R., Van Belleghem, S. M., Wright, D. S., Kuo, C.-Y., Lozano-Urrego, D., & Maulana, A. (2024). Adaptive introgression of a visual preference gene. Science, 383(6689), 1368–1373.

Rubin, C.-J., Enbody, E. D., Dobreva, M. P., Abzhanov, A., Davis, B. W., Lamichhaney, S., Pettersson, M., Sendell-Price, A. T., Sprehn, C. G., & Valle, C. A. (2022). Rapid adaptive radiation of Darwin’s finches depends on ancestral genetic modules. Science Advances, 8(27), eabm5982.

Schluter, D. (1996). Adaptive radiation along genetic lines of least resistance. Evolution, 50(5), 1766–1774.

Short, A. W., & Streisfeld, M. A. (2023). Ancient hybridization leads to the repeated evolution of red flowers across a monkeyflower radiation. Evolution letters. 10.1093/evlett/qrad024

Sidlauskas, B. (2008). Continuous and arrested morphological diversification in sister clades of characiform fishes: a phylomorphospace approach. Evolution, 62(12), 3135–3156.

Sinnott-Armstrong, M. A., Deanna, R., Pretz, C., Liu, S., Harris, J. C., Dunbar-Wallis, A., Smith, S. D., & Wheeler, L. C. (2022). How to approach the study of syndromes in macroevolution and ecology. Ecology and evolution, 12(3), e8583.

Šlenker, M., Koutecký, P., & Marhold, K. (2022). MorphoTools2: an R package for multivariate morphometric analysis. Bioinformatics, 38(10), 2954–2955.

Smith, S. A., & O’Meara, B. C. (2012). treePL: divergence time estimation using penalized likelihood for large phylogenies. Bioinformatics, 28(20), 2689–2690.

Stayton, C. T. (2015). The definition, recognition, and interpretation of convergent evolution, and two new measures for quantifying and assessing the significance of convergence. Evolution, 69(8), 2140–2153.

Stebbins, G. (1989). Adaptive shifts toward hummingbird pollination. The evolutionary ecology of plants, 39–60.

Stebbins, G. L. (1970). Adaptive radiation of reproductive characteristics in angiosperms, I: pollination mechanisms. Annual Review of Ecology and Systematics, 1(1), 307–326.

Stebbins, G. L. (1974). Flowering plants: evolution above the species level. Belknap Press.

Stoltzfus, A., & McCandlish, D. M. (2017). Mutational biases influence parallel adaptation. Molecular biology and evolution, 34(9), 2163–2172.

Stone, B. W., & Wessinger, C. A. (2024). Ecological diversification in an adaptive radiation of plants: the role of de novo mutation and introgression. Molecular biology and evolution, 41(1), msae007.

Stone, B. W., & Wolfe, A. D. (2021). Asynchronous rates of lineage, phenotype, and niche diversification in a continental-scale adaptive radiation. bioRxiv. 10.1101/2021.06.14.448393

Suvorov, A., Kim, B. Y., Wang, J., Armstrong, E. E., Peede, D., D’agostino, E. R., Price, D. K., Waddell, P. J., Lang, M., & Courtier-Orgogozo, V. (2022). Widespread introgression across a phylogeny of 155 Drosophila genomes. Current Biology, 32(1), 111–123. e115.

Swift, J., Luginbuehl, L. H., Hua, L., Schreier, T. B., Donald, R. M., Stanley, S., Wang, N., Lee, T. A., Nery, J. R., & Ecker, J. R. (2024). Exaptation of ancestral cell-identity networks enables C4 photosynthesis. Nature, 636(8041), 143–150.

Thomson, J. D., & Wilson, P. (2008). Explaining evolutionary shifts between bee and hummingbird pollination: convergence, divergence, and directionality. International Journal of Plant Sciences, 169(1), 23–38.

Wessinger, C. A., Freeman, C. C., Mort, M. E., Rausher, M. D., & Hileman, L. C. (2016). Multiplexed shotgun genotyping resolves species relationships within the North American genus Penstemon. American Journal of Botany, 103(5), 912–922.

Wessinger, C. A., & Hileman, L. C. (2016). Accessibility, constraint, and repetition in adaptive floral evolution. Developmental biology, 419(1), 175–183.

Wessinger, C. A., Hileman, L. C., & Rausher, M. D. (2014). Identification of major quantitative trait loci underlying floral pollination syndrome divergence in *Penstemon*. Phil. Trans. R. Soc. B, 369(1648), 20130349.

Wessinger, C. A., Katzer, A. M., Hime, P. M., Rausher, M. D., Kelly, J. K., & Hileman, L. C. (2023). A few essential genetic loci distinguish Penstemon species with flowers adapted to pollination by bees or hummingbirds. PLoS Biology, 21(9), e3002294.

Wessinger, C. A., & Kelly, J. K. (2018). Selfing can facilitate transitions between pollination syndromes. The American Naturalist, 191(5), 582–594.

Wessinger, C. A., Kelly, J. K., Jiang, P., Rausher, M. D., & Hileman, L. C. (2018). SNP-skimming: A fast approach to map loci generating quantitative variation in natural populations. Molecular ecology resources, 18(6), 1402–1414.

Wessinger, C. A., & Rausher, M. D. (2015). Ecological transition predictably associated with gene degeneration. Molecular biology and evolution, 32(2), 347–354.

Wilson, P., Castellanos, M. C., Hogue, J. N., Thomson, J. D., & Armbruster, W. S. (2004). A multivariate search for pollination syndromes among penstemons. Oikos, 104(2), 345–361.

Wilson, P., Castellanos, M. C., Wolfe, A. D., & Thomson, J. D. (2006). Shifts between bee and bird pollination in Penstemons. Plant–pollinator interactions: from specialization to generalization, 47–68.

Wilson, P., & Jordan, E. A. (2009). Hybrid intermediacy between pollination syndromes in Penstemon, and the role of nectar in affecting hummingbird visitation. Botany, 87(3), 272–282.

Wilson, P., & Valenzuela, M. (2002). Three naturally occurring Penstemon hybrids. Western North American Naturalist, 25–31.

Wilson, P., Wolfe, A. D., Armbruster, W. S., & Thomson, J. D. (2007). Constrained lability in floral evolution: counting convergent origins of hummingbird pollination in Penstemon and Keckiella. New Phytologist, 176(4), 883–890.

Wogan, G. O., Yuan, M. L., Mahler, D. L., & Wang, I. J. (2023). Hybridization and transgressive evolution generate diversity in an adaptive radiation of Anolis lizards. Systematic biology, 72(4), 874–884.

Wolfe, A. D., Blischak, P. D., & Kubatko, L. (2021). Phylogenetics of a rapid, continental radiation: diversification, biogeography, and circumscription of the beardtongues (Penstemon; Plantaginaceae). bioRxiv. 10.1101/2021.04.20.440652

Wolfe, A. D., Randle, C. P., Datwyler, S. L., Morawetz, J. J., Arguedas, N., & Diaz, J. (2006). Phylogeny, taxonomic affinities, and biogeography of Penstemon (Plantaginaceae) based on ITS and cpDNA sequence data. American Journal of Botany, 93(11), 1699–1713.

Zhang, C., Rabiee, M., Sayyari, E., & Mirarab, S. (2018). ASTRAL-III: polynomial time species tree reconstruction from partially resolved gene trees. BMC bioinformatics, 19(6), 153.

