## Supplemental Material for "Rapid floral syndrome convergence in *Penstemon* through independent genetic variation"

### Supplemental Materials and Methods

#### Phenotyping

We measured the following floral dimension traits on whole flowers using digital calipers: floral tube length (length from the base of the flower to the tube opening), tube width (width at the tube opening), long stamen length (length from the base of the flower to the end of the lateral stamens), short stamen length (length from the base of the flower to the end of the ventral stamens), and style length (length from the base of the flower to the end of the pistil). We measured nectar volume in microliters from flowers on the first day of opening using 5 μl micro-capillary tubes (Drummond Scientific, Broomali, PA, USA). We imaged the nectaries found at the base of the short stamen filaments alongside a millimeter ruler under a Dino-Lite Premier digital microscope (Dino-Lite, Torrance, CA, USA) set to 20x magnification and measured nectary area from the resulting photographs using *FIJI* (Schindelin et al., 2012).

We extracted anthocyanidin pigments from pressed floral tissue using an isoamyl alcohol extraction protocol (Harborne, 1984) and separated pigments using thin layer chromatography on Cellulose F glass plates (Millipore, Burlington, MA, USA), following Stevens et al. (2023). We compared extracted pigments to the standard anthocyanidins pelargonidin, cyanidin, and delphinidin and scored for presence of each type. Note that methylated anthocyanidin pigments are not produced by *Penstemon* species (Wessinger & Rausher, 2015). We assigned each accession a numeric value based on their qualitative anthocyanidin content that reflects the number of hydroxyl groups on the central B-ring of the anthocyanidin pigment: 1=pelargonidin, 2=cyanidin, 3=delphinidin. Individuals producing a combination of pigments were assigned the average of the two values (e.g., 2.5 assigned to individuals producing cyanidin and delphinidin).

We used reflectance spectroscopy to measure the flower color of the upper petal lobe surface with a UV-vis spectrometer (FLAME-S-UV-VIS, Ocean Insight, Dunedin, FL, USA), equipped with a Y-shaped reflection probe (R400-7-UV-VIS, Ocean Insight) that supplied light from a deuterium/tungsten light source (DH-200-BAL, Ocean Insight). We calibrated reflectance measurements with a Spectralon diffuse reflectance standard (WS-1, Ocean Insight) before recording reflectance data from 300 to 800 nm using OceanView software (Ocean Insight) in a darkened room with no overhead lighting. We sandwiched petal tissue between two clean glass microscope slides to eliminate spectral variation due to petal curvature and then measured reflectance with the reflectance probe at a 45-degree angle relative to the sample. We performed these measurements on 25 of the 34 sampled individuals.

We processed raw reflectance spectra using the R package *pavo* (Maia et al., 2019). We calculated hue and chroma using segment classification (Smith, 2014). Hue is calculated in degrees – a circular metric – and because the values in our dataset bridge the 0°/360° boundary, we calculated an "adjusted hue" value by by adding 90° to each hue value. We also estimated the detectability of each floral reflectance spectrum to bees vs. hummingbirds by comparing chromatic contrast values under a honeybee visual model vs. an average avian violet sensitive model (to represent hummingbird vision) using *pavo.* We calculated quantum catch for each photoreceptor against a background spectrum measured from a *Penstemon* leaf using the *pavo* function *vismodel*. We then used the *pavo* function *coldist* to calculate distances between colors in each pollinators' visual space under a receptor-noise model (Vorobyev & Osorio, 1998). This model yields chromatic contrast values in units of Just Noticeable Differences (JNDs), where values below 1 are assumed not to be visually detectable and increasing JND values indicate the color is increasingly detectable.

#### Whole genome resequencing

We extracted DNA from fresh or silica-dried leaf tissue using a modified CTAB protocol. Whole genome Illumina library preparations were generated using the Illumina DNA Prep kit and resulting libraries were sequenced to ~5x depth using 150-bp paired-end reads generated on the NovaSeq X platform (Illumina, San Diego, CA, USA) at the Duke University Sequencing and Genomic Technologies research core facility.

Using *fastp* version 0.23.2 (Chen et al., 2018), we trimmed adapters and poly-x on the 3'-ends of reads, corrected bases for overlapping reads, and filtered out unpaired reads or those less than 30 bp. We then checked filtered sequence quality using *fastqc* version 0.11.8 (Andrews et al., 2010). We mapped our filtered reads to the annotated *P. eatonii* reference genome (Jarvis et al., 2025) using *bwa mem* version 0.7.17 (Li, 2013), marked and removed duplicated reads using *samtools* version 1.51.1 (Li et al., 2009), and clipped overlapping paired-end reads using *bamutil* clipOverlap (Jun et al., 2015).

We generated an all sites vcf file using *bcftools* (Li, 2011) in single-sample calling mode, using a minimum mapping quality of 30, a minimum base quality of 20, and excluding sites within 5 bp of an indel. We excluded sites within annotated repeats using *bcftools* and retained only those sites that mapped to the 8 major chromosomes of the reference genome. As a final filtering step, we filtered on minimum genotype call quality (2), maximum number of alleles (2), minimum read depth (2), maximum mean depth (16), maximum per-site proportion of heterozygote calls (75%), and maximum per-site proportion of missing individuals (10%) using *vcftools* version 0.1.17 (Danecek et al., 2011).

**Tree inference**

We generated whole-genome consensus sequences using the *consensus* function of *bcftools*. We then generated two sets of sequence alignments: 10 kb non-overlapping windows and coding sequences (CDS) using custom python scripts reported in Stone and Wessinger (2024). We then inferred trees for each sequence alignment using *IQ-TREE* version 2.1.2 (Nguyen et al., 2015) under a general time reversal model, allowing invariable sites and rate variation across sites (-m GTR+I+G). We used *ASTRAL-III* version 5.7.1 (Zhang et al., 2018) to infer a CDS-based species tree from the collection of CDS trees and a 10 kb window-based species tree from the collection of 10 kb window trees. We rooted each tree using the outgroup taxon *P. palmeri*. We calculated gene concordance factors on each species tree using *IQ-TREE* (Minh et al., 2020).

We inferred relative divergence times for the 10 kb window-based ASTRAL tree using a penalized likelihood approach implemented in *treePL* version 1.0 (Smith & O’Meara, 2012). Specifically, we first used *treePL* to test optimal settings using the 'prime' mode and then ran a final analysis using the 'thorough' mode and the following program settings: opt=5, optad=5, and optcvad=2. We used this ultrametric tree for all phylogenetic comparative analyses.

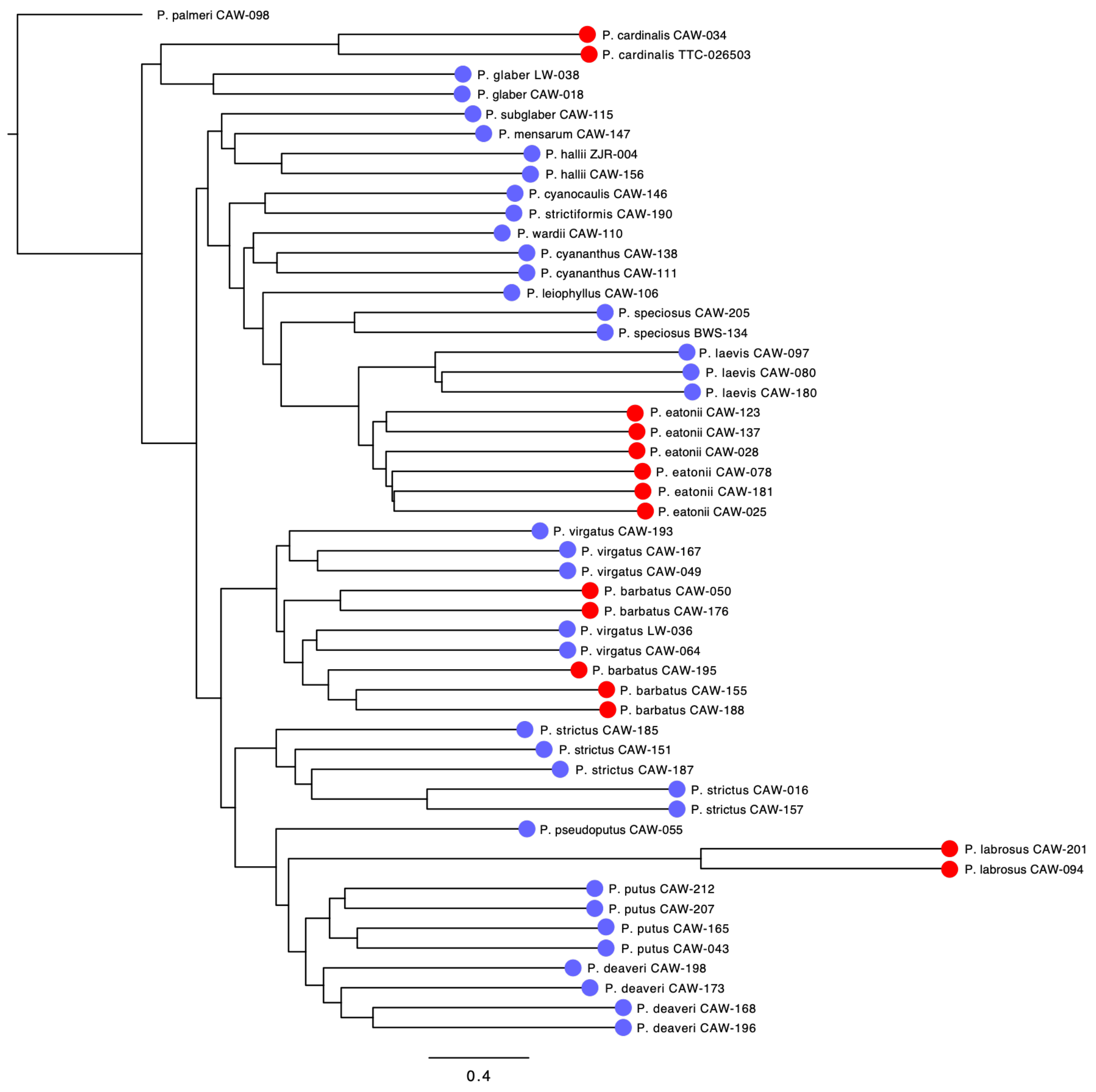


Figure S1. Phylogenetic relationships between accessions inferred using a species tree approach based on CDS regions. Purple: bee syndrome species; red: hummingbird syndrome species.

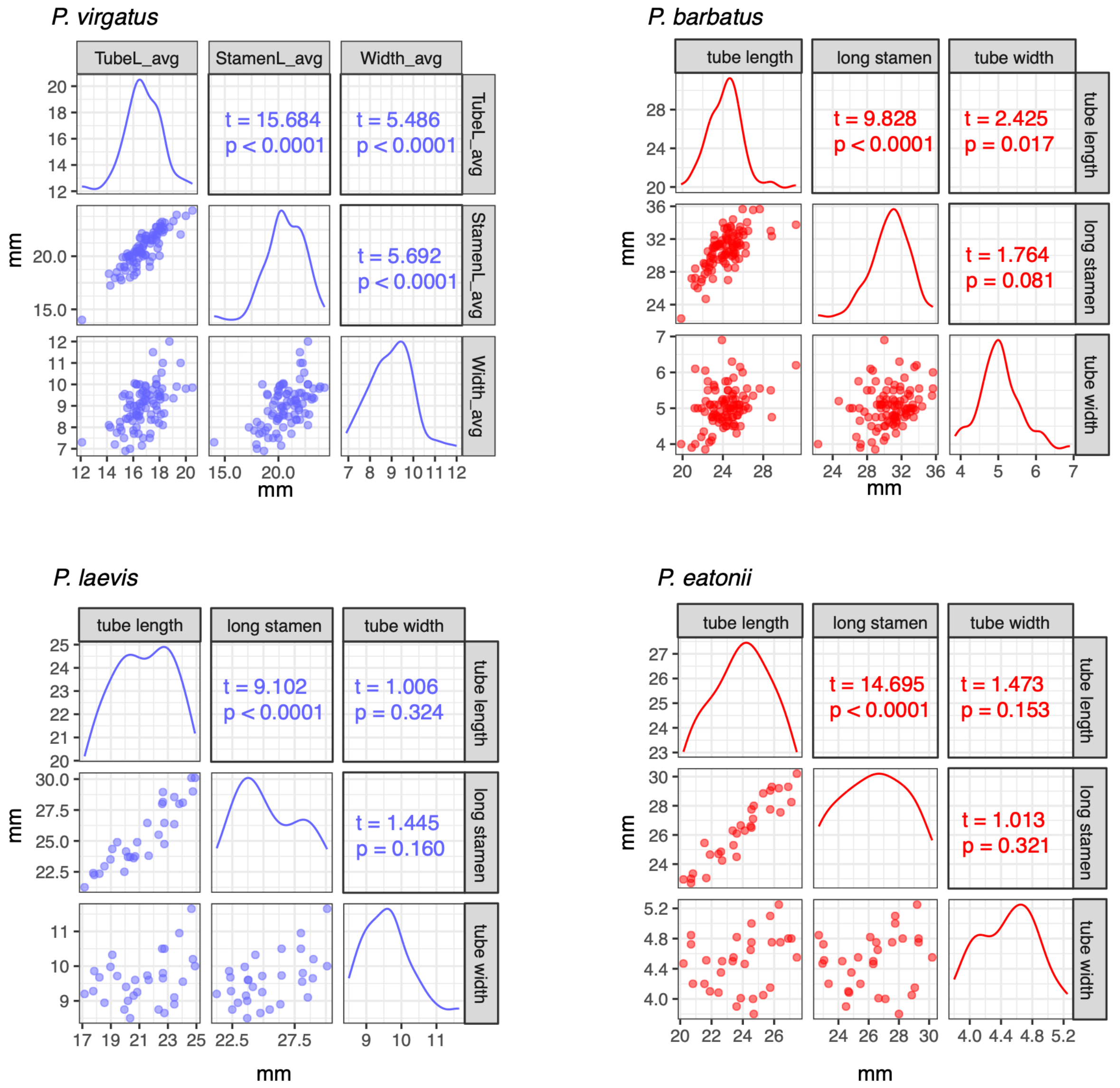


Figure S2. Relationships among phenotypes measured with populations of two bee syndrome species (*P. virgatus* and *P. laevis*) and two hummingbird syndrome species (*P. barbatus* and *P. eatonii*). Reported are t- and p-values from linear models described in the main text.

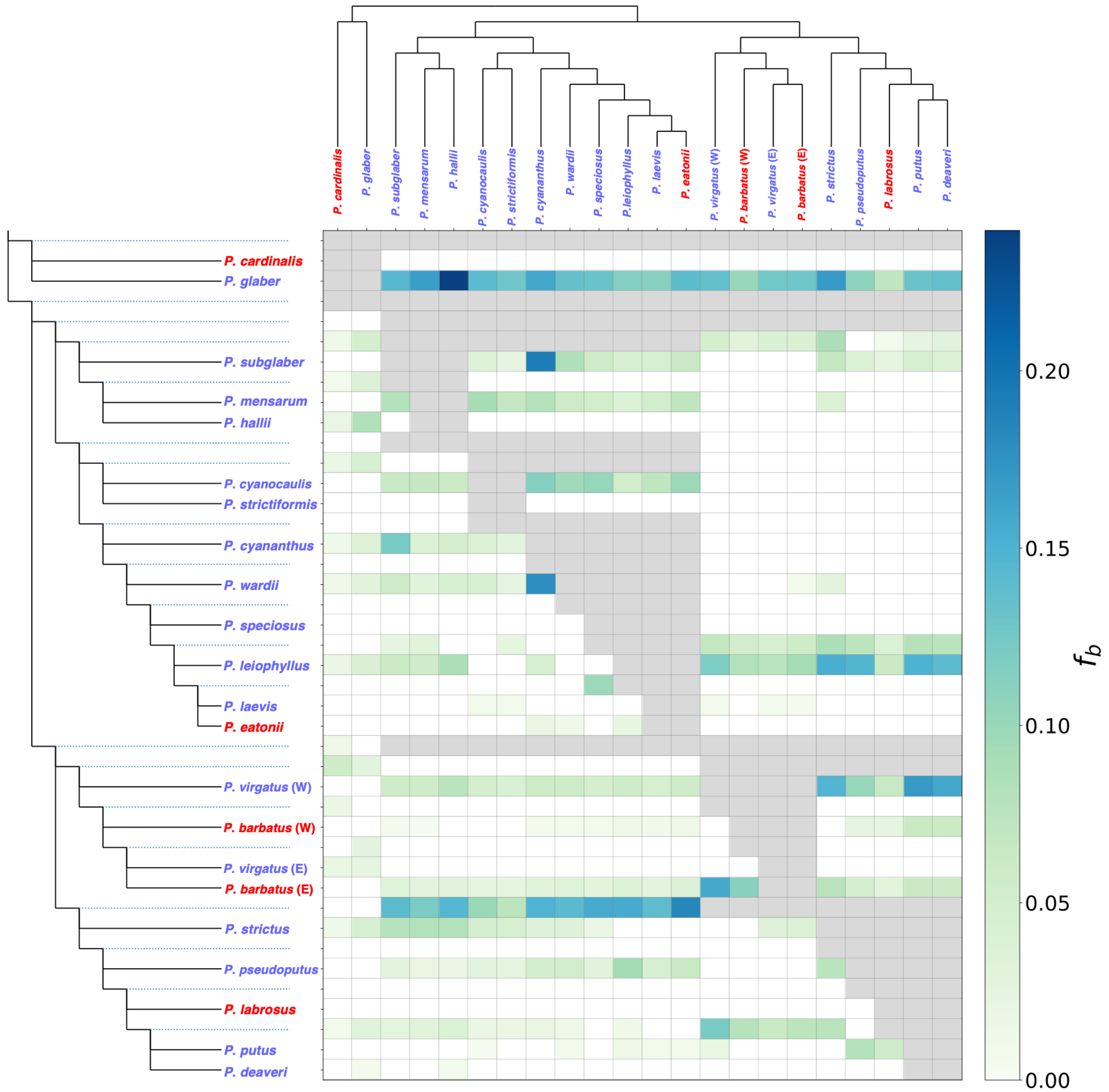


Figure S3. Results of *f*-branch analysis testing for excess allele sharing between branches and tips in the species tree, which is interpreted as evidence for introgression between species and/or clades. The species tree used here is the ASTRAL tree based on 10 kb genomic windows. Red: hummingbird syndrome species, purple: bee syndrome species.

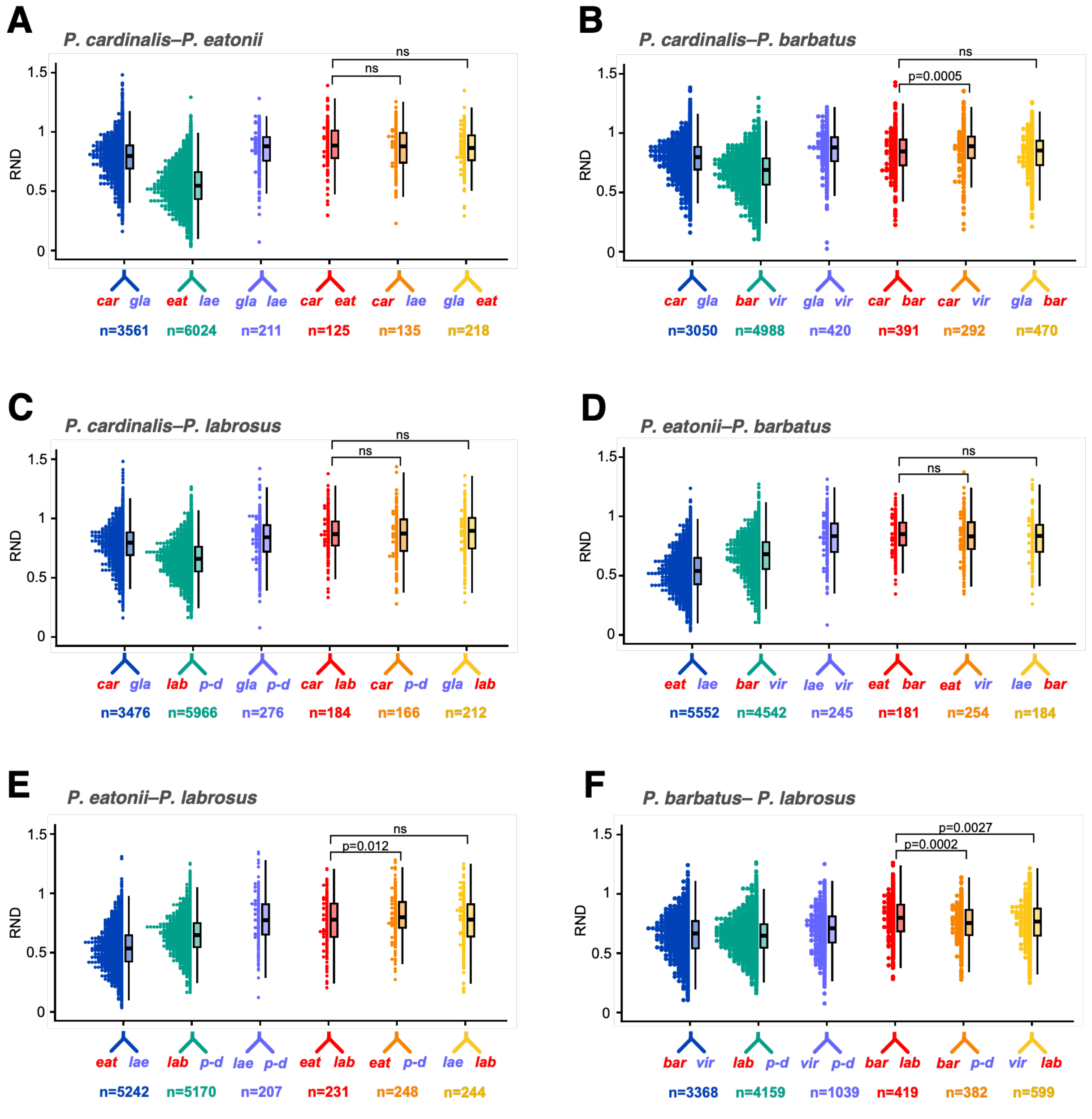


Figure S4. Distributions of relative node depth (RND) for 10 kb regions of the genome showing allele sharing between the following taxa: H1-B1 (navy), H2-B2 (green), B1-B2 (purple), H1-H2 (red), H1-B2 (orange), and H2-B1 (yellow). Significance of Mann-Whitney U tests are reported. Ns: $p > 0.05$.

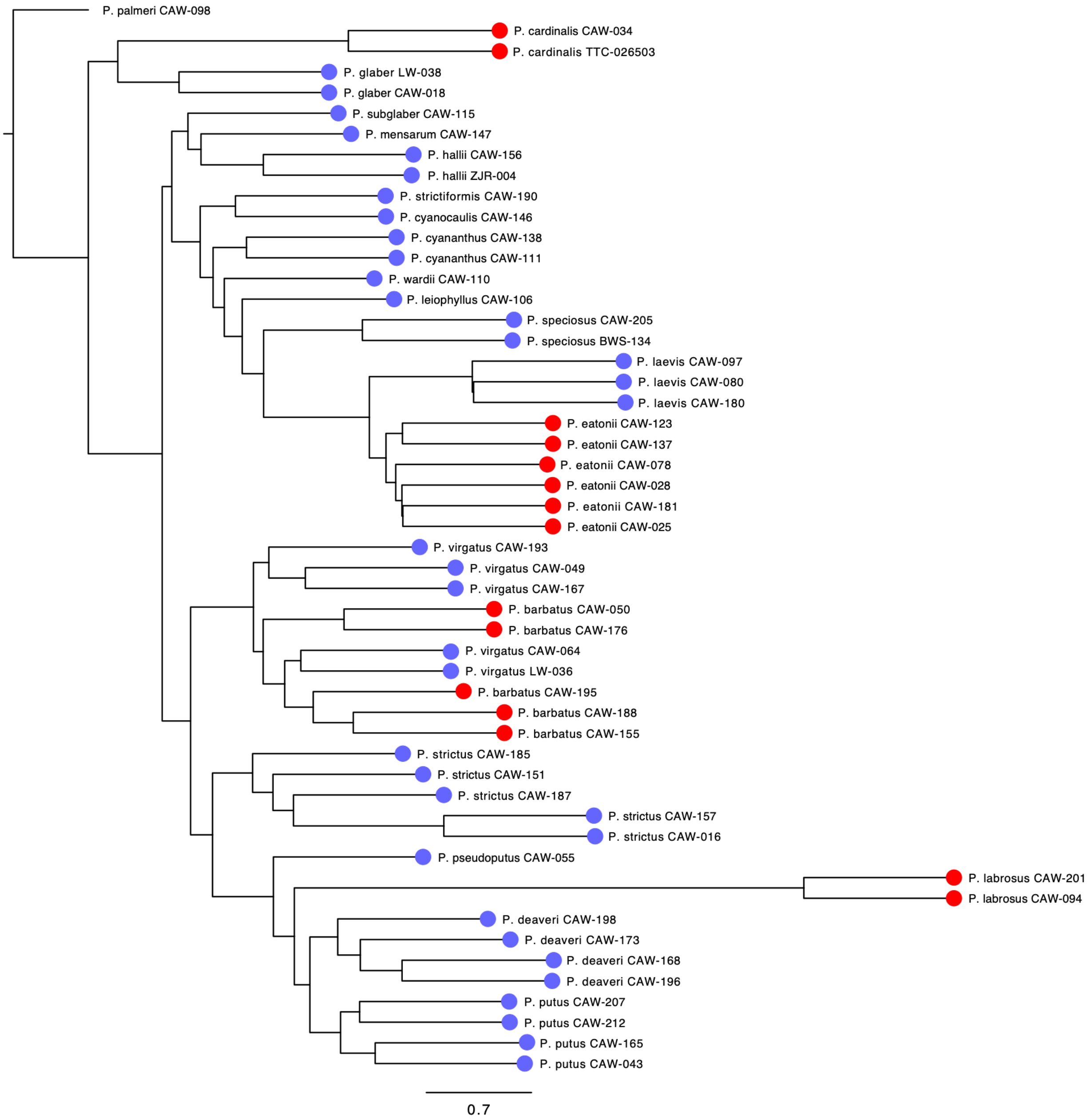


Figure S5. Phylogenetic relationships between accessions inferred using a species tree approach based on 10 kb window regions and using the *P. barbatus* reference genome. Purple: bee syndrome species; red: hummingbird syndrome species.

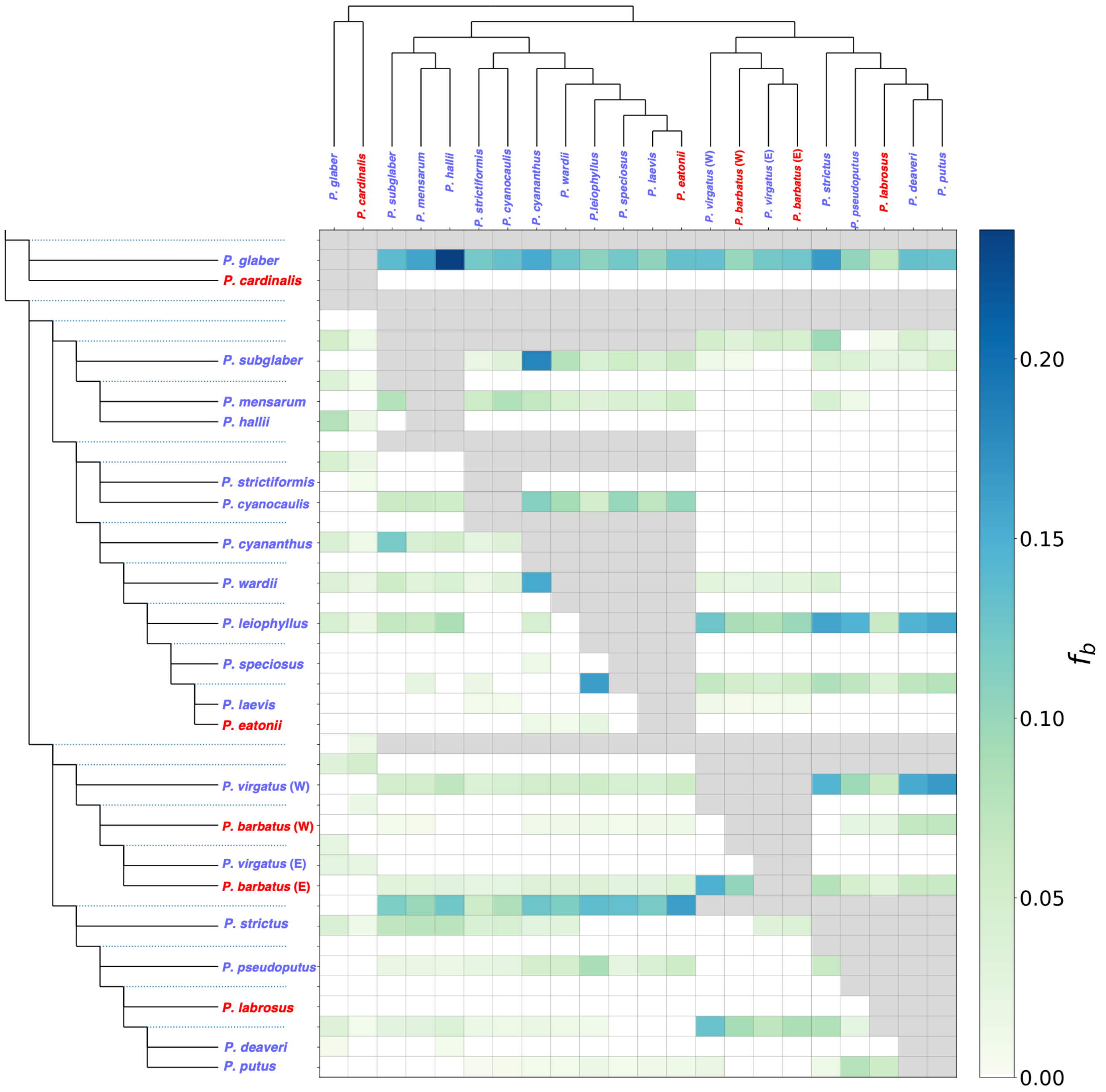


Figure S6. Results of *f*-branch analysis testing for excess allele sharing between branches and tips in the species tree using the *P. barbatus* reference genome.The species tree used here is the ASTRAL tree based on 10 kb genomic windows. Red: hummingbird syndrome species, purple: bee syndrome species.

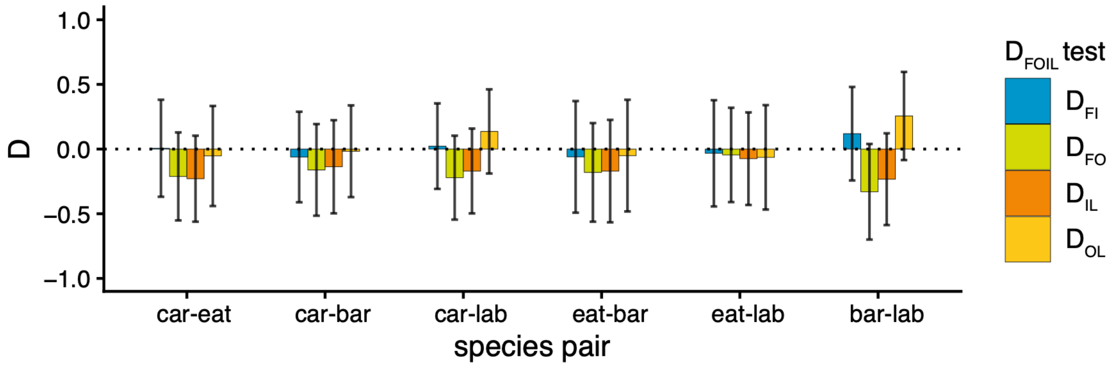


Figure S7. Results of $D_{FOIL}$ tests for all six pairwise tests of introgression, using the *P. barbatus* reference genome.

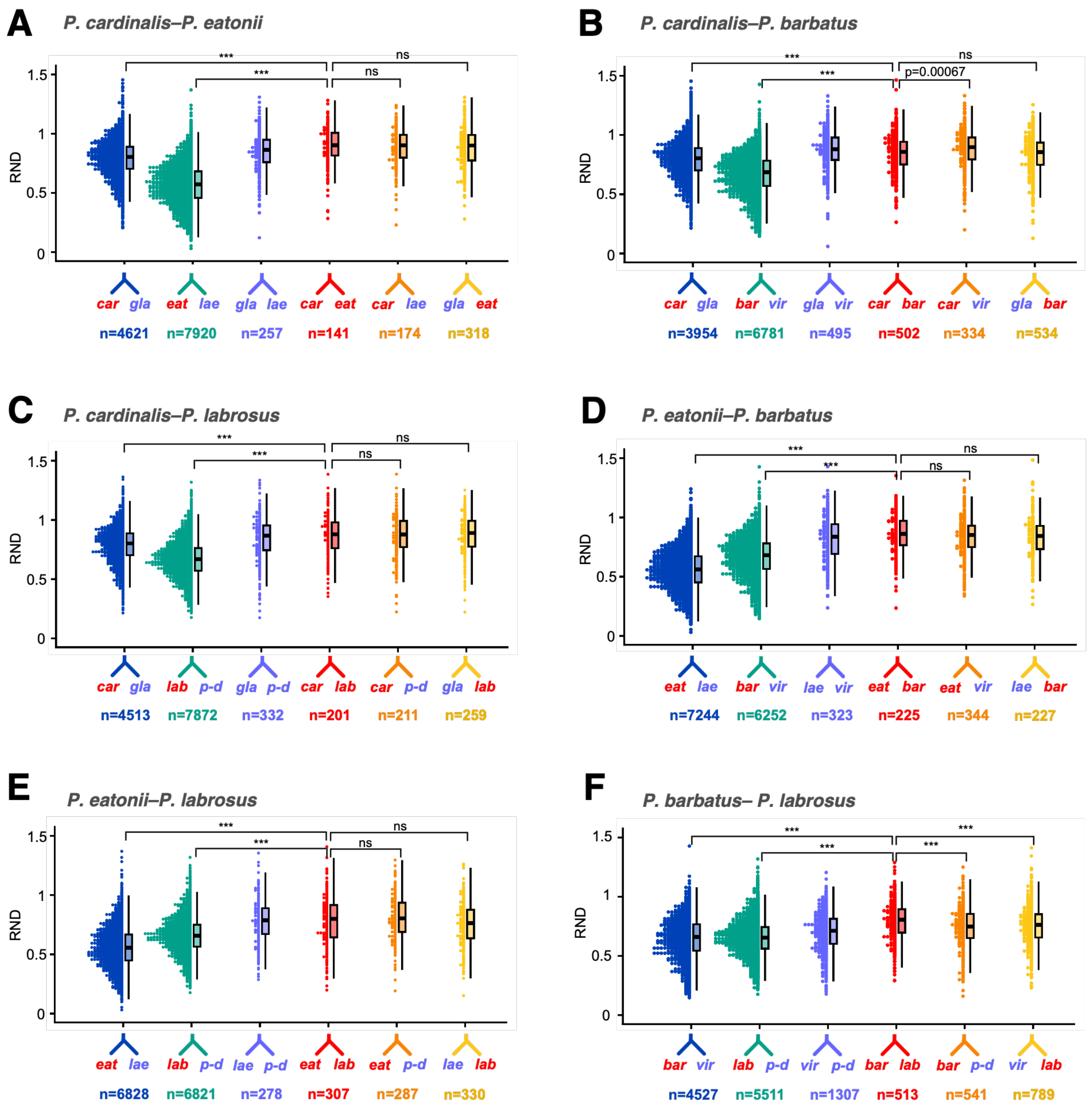


Figure S8. Distributions of relative node depth (RND) for 10 kb regions of the genome (*P. barbatus* reference genome) showing allele sharing between the following taxa: H1-B1 (navy), H2-B2 (green), B1-B2 (purple), H1-H2 (red), H1-B2 (orange), and H2-B1 (yellow). Significance of Mann-Whitney U tests are reported. Ns: $p > 0.05$, ***: $p < 0.0001$.

Table S1. Collection information for *Penstemon* samples used in this study. BWS: Benjamin Stone, CAW: Carolyn Wessinger; COLO: University of Colorado Museum of Natural History; KANU: R.L. McGregor Herbarium at University of Kansas; LCW: Lucas Wheeler; MJ: Matthew Johnson; TTC: E.L. Reed Herbarium at Texas Tech University; ZJR: Zachary Radford. Asterisks indicate accessions used in 5-taxon tests for introgression.

| **Population identifier** | **Species determination (abbreviation in Figure 1)** | **Locality of population** | **Collector** | **Herbarium deposited** | **Phenotyped** |
|---|---|---|---|---|---|
| BWS-134 | *P. speciosus* | Mono Pass, CA | BWS | KANU | no |
| CAW-016 | P. strictus (str-016) | Victor, CO | CAW | KANU | yes |
| CAW-018* | *P. glaber* | Midland, CO | CAW | KANU | no |
| CAW-025 | *P. eatonii* | Cortez, CO | CAW | KANU | no |
| CAW-028 | *P. eatonii* (eat-028) | McDowell, AZ | CAW | KANU | yes |
| CAW-034* | *P. cardinalis* (car-034) | Bonito Lake, NM | CAW | KANU | yes |
| CAW-043* | *P. putus* | Pine, AZ | CAW | KANU | no |
| CAW-049* | *P. virgatus* (vir-049) | Kendrick Park, AZ | CAW | KANU | yes |
| CAW-050 | *P. barbatus* (bar-050) | Flagstaff, AZ | CAW | KANU | yes |
| CAW-055 | *P. pseudoputus* (psp-055) | Kaibab Plateau, AZ | CAW | KANU | yes |
| CAW-064 | *P. virgatus* (vir-064) | Cloudcroft, NM | CAW | KANU | yes |
| CAW-078 | *P. eatonii* | Mt. Carmel Jct., UT | CAW | KANU | no |
| CAW-080* | *P. laevis* | Mt. Carmel Jct., UT | CAW | KANU | no |
| CAW-094* | *P. labrosus* (lab-094) | Big Bear Lake, CA | CAW | KANU | yes |
| CAW-097 | *P. laevis* (lae-097) | Rosy Canyon, AZ | CAW | KANU | yes |
| CAW-098* | *P. palmeri* | Rosy Canyon, AZ | CAW | KANU | no |
| CAW-106 | *P. leiophyllus* (lei-106) | Cedar Canyon, UT | CAW | KANU | yes |
| CAW-110 | *P. wardii* | Soldier Canyon, UT | CAW | KANU | no |
| CAW-111 | *P. cyananthus* | Nebo Loop, UT | CAW | KANU | no |
| CAW-115 | *P. subglaber* (sub-115) | Tucker, UT | CAW | KANU | yes |
| CAW-123* | *P. eatonii* | Antimony, UT | CAW | KANU | no |
| CAW-137 | *P. eatonii* | Three Creeks, UT | CAW | KANU | no |
| CAW-138 | *P. cyananthus* | Three Creeks, UT | CAW | KANU | no |
| CAW-146 | *P. cyanocaulis* | Devil's Kitchen, CO | CAW | KANU | no |
| CAW-147 | *P. mensarum* (men-147) | Powderhorn Ski, CO | CAW | KANU | yes |
| CAW-151 | *P. comarrhenus* | Mesa Creek, CO | CAW | KANU | no |
| CAW-155* | *P. barbatus* | Cottonwood Pass, CO | CAW | KANU | no |
| CAW-156 | *P. hallii* | Fremont Pass, CO | CAW | KANU | no |
| CAW-157 | *P. strictus* (str-157) | Keystone, CO | CAW | KANU | yes |
| CAW-165 | *P. putus* (put-165) | Forest Lakes, AZ | CAW | KANU | yes |
| CAW-167 | *P. virgatus* (vir-167) | Overgaard, AZ | CAW | KANU | yes |
| CAW-168 | *P. deaveri* | Sheeps Crossing, AZ | CAW | KANU | no |
| CAW-173 | *P. deaveri* (dea-173) | Willow Creek, NM | CAW | KANU | yes |
| CAW-176 | *P. barbatus* (bar-176) | Pinos Altos, NM | CAW | KANU | yes |
| CAW-180 | *P. laevis* | Rosy Canyon, UT | CAW | KANU | no |
| CAW-181 | *P. eatonii* (eat-181) | Gooseberry Mesa, UT | CAW | KANU | yes |
| CAW-185 | *P. strictus* | Eagles Nest Lake, NM | CAW | KANU | no |
| CAW-187 | *P. strictus* (str-187) | Horca, CO | CAW | KANU | yes |
| CAW-188 | *P. barbatus* (bar-188) | La Manga Pass, CO | CAW | KANU | yes |
| CAW-190 | *P. strictiformis* (stf-190) | Llaves, NM | CAW | KANU | yes |
| CAW-193 | *P. virgatus* (vir-193) | Mt. Taylor, NM | CAW | KANU | yes |
| CAW-195 | *P. barbatus* (bar-195) | Zuni Canyon, NM | CAW | KANU | yes |
| CAW-196 | *P. deaveri* (dea-196) | Gallo Peak, NM | CAW | KANU | yes |
| CAW-198 | *P. deaveri* (dea-198) | Taylor Peak, NM | CAW | KANU | yes |
| CAW-201 | *P. labrosus* (lab-201) | Big Bear Lake, CA | CAW | KANU | yes |
| CAW-205 | *P. speciosus* (spe-205) | Sunriver, OR | CAW | KANU | yes |
| CAW-207 | *P. putus* (vir-207) | Humphreys Peak, AZ | CAW | KANU | yes |
| CAW-212 | *P. putus* | Sycamore Falls, AZ | CAW | KANU | no |
| LW-036 | *P. virgatus* | Four Mile Canyon, CO | LW | COLO | no |
| LW-038 | *P. glaber* (gla-038) | Brainard Lake, CO | LW | COLO | yes |
| TTC026503 | *P. cardinalis* | Guadalupe Mtns, TX | MJ | TTC | no |
| ZJR-004 | *P. hallii* (hal-004) | Guanella Pass, CO | ZJR | KANU | yes |

Table S2. Information for populations phenotyped for floral dimension traits in the field. CAW: Carolyn Wessinger; KANU: R.L. McGregor Herbarium at University of Kansas.

| **Population** | **Species** | **Locality** | **Collector** | **Herbarium deposited** |
|---|---|---|---|---|
| CAW015 | *P. virgatus* | Victor, CO | CAW | KANU |
| CAW017 | *P. virgatus* | Lake George, CO | CAW | KANU |
| CAW019 | *P. virgatus* | Buena Vista, CO | CAW | KANU |
| CAW020 | *P. virgatus* | Leadville, CO | CAW | KANU |
| CAW030 | *P. barbatus* | Gila NF, NM | CAW | KANU |
| CAW031 | *P. virgatus* | Cloudcroft, NM | CAW | KANU |
| CAW032 | *P. virgatus* | Ski Cloudcroft, NM | CAW | KANU |
| CAW033 | *P. virgatus* | Alto, NM | CAW | KANU |
| CAW035 | *P. virgatus* | Corona, NM | CAW | KANU |
| CAW036 | *P. barbatus* | Valle Caldera, NM | CAW | KANU |
| CAW037 | *P. barbatus* | Apache Canyon, NM | CAW | KANU |
| CAW041 | *P. barbatus* | Questa, NM | CAW | KANU |
| CAW080 | *P. laevis* | Mt. Carmel Jct, UT | CAW | KANU |
| CAW162 | *P. barbatus* | Lake Mary, AZ | CAW | KANU |
| CAW164 | *P. barbatus* | Mogollon Rim, AZ | CAW | KANU |
| CAW167 | *P. virgatus* | Overgaard, AZ | CAW | KANU |
| CAW169 | *P. barbatus* | Greer, AZ | CAW | KANU |
| CAW174 | *P. barbatus* | Mogollon, NM | CAW | KANU |
| CAW176 | *P. barbatus* | Pinos Altos, NM | CAW | KANU |
| CAW177 | *P. barbatus* | High Rolls, NM | CAW | KANU |
| CAW180 | *P. laevis* | Rosy Canyon, AZ | CAW | KANU |
| CAW181 | *P. eatonii* | Gooseberry Mesa, UT | CAW | KANU |
| CAW232 | *P. eatonii* | Kolob Canyon, UT | CAW | KANU |
| CAW233 | *P. laevis* | Red Hollow Canyon, UT | CAW | KANU |
| CAW234 | *P. eatonii* | Little Creek Mesa, UT | CAW | KANU |

Table S3. Results of phylogenetic ANOVA models implemented in *caper* testing the effect of pollination syndrome on floral trait variation. Lambda refers to the estimated Pagel's lambda parameter.

| **Model** | **lambda** | **DF** | **F-value** | **p-value** |
|---|---|---|---|---|
| long stamen length ~ syndrome | 0 | 28 | 53.949 | 5.353 e -08 |
| short stamen length ~ syndrome | 0.54 | 28 | 65.931 | 7.696 e -09 |
| style length ~ syndrome | 0.48 | 28 | 44.726 | 2.937 e -07 |
| tube length ~ syndrome | 0 | 28 | 59.01 | 2.282 e -08 |
| tube width ~ syndrome | 0.70 | 28 | 49.212 | 1.25 e -07 |
| log nectar volume ~ syndrome | 0 | 28 | 31.35 | 5.411 e -06 |
| log nectary area ~ syndrome | 1 | 28 | 58.258 | 2.582 e -08 |
| length/width ratio ~ syndrome | 0.16 | 28 | 302.04 | < 2.2 e -16 |
| anthocyanin ~ syndrome | 1 | 28 | 619.57 | < 2.2 e -16 |
| PC1 ~ syndrome | 0.46 | 28 | 141.76 | 1.792 e -12 |
| PC2 ~ syndrome | 0.44 | 28 | 3.0033 | 0.0941 |
| adjusted hue ~ syndrome | 0 | 23 | 309.61 | 7.73 e -15 |
| chroma ~ syndrome | 0 | 23 | 153.68 | 9.475 e -12 |
| brightness ~ syndrome | 0 | 23 | 15.032 | 0.000763 |
| chromatic contrast bee ~ syndrome | 0 | 23 | 57.507 | 1.061 e -07 |
| chromatic contrast bird ~ syndrome | 0 | 23 | 112.83 | 2.42 e -10 |

Table S4. Results of phylogenetic ANOVA models implemented in *caper* testing whether floral traits in bee syndrome species differ according to whether or not the accession is a sister taxon to hummingbird syndrome species. Lambda refers to the estimated Pagel's lambda parameter.

| **Model** | **DF** | **lambda** | **F-value** | **p-value** |
|---|---|---|---|---|
| long stamen length ~ sister status | 19 | 1 | 2.2707 | 0.1483 |
| short stamen length ~ sister status | 19 | 1 | 3.713 | 0.0691 |
| style length ~ sister status | 19 | 1 | 2.4001 | 0.1378 |
| tube length ~ sister status | 19 | 0 | 0.4426 | 0.5139 |
| tube width ~ sister status | 19 | 1 | 0.8223 | 0.3759 |
| log nectar volume ~ sister status | 19 | 0 | 1.9712 | 0.1765 |
| log nectary area ~ sister status | 19 | 0 | 0.4395 | 0.5153 |
| length / width ratio ~ sister status | 19 | 0 | 1.7317 | 0.2038 |
| anthocyanin ~ sister status | 19 | 1 | 2.521 | 0.1288 |
| PC1 ~ sister status | 19 | 0.6 | 2.7666 | 0.1127 |
| PC2 ~ sister status | 19 | 0 | 2.4233 | 0.136 |

Table S5. Phylogenetic correlations among traits in bee syndrome species estimated in *caper*.

| **Model** | **Correlation coefficient** | **p-value** |
|---|---|---|
| Long stamen length, short stamen length | 0.9684 | 6.403 e -13 |
| Long stamen length, style length | 0.8671 | 3.634 e -07 |
| Long stamen length, tube length | 0.8608 | 5.051 e -07 |
| Long stamen length, tube width | 0.2111 | 0.3584 |
| Long stamen length, log nectar volume | 0.5232 | 0.0149 |
| Long stamen length, long nectary area | 0.4570 | 0.0373 |
| Short stamen length, style length | 0.8481 | 1.201 e -06 |
| Short stamen length, tube length | 0.8621 | 5.065 e -07 |
| Short stamen length, tube width | 0.2983 | 0.189 |
| Short stamen length, log nectar volume | 0.5621 | 0.0080 |
| Short stamen length, log nectary area | 0.4211 | 0.0573 |
| Style length, tube length | 0.8141 | 7.094 e -06 |
| Style length, tube width | 0.2427 | 0.2892 |
| Style length, log nectar volume | 0.6149 | 0.0030 |
| Style length, log nectary area | 0.3457 | 0.1248 |
| Tube length, tube width | 0.3140 | 0.1573 |
| Tube length, log nectar volume | 0.5705 | 0.0069 |
| Tube length, log nectary area | 0.3894 | 0.0810 |
| Tube width, log nectar volume | 0.4189 | 0.0587 |
| Tube width, log nectary area | 0.3564 | 0.1128 |
| Log nectar volume, log nectary area | 0.5406 | 0.0114 |

Table S6. Statistical comparisons of RND distributions between taxa in a 5-taxon testing framework. Reference genome used for analyses was *P. eatonii*.

| **B1 taxon** | **H1 taxon** | **H2 taxon** | **B2 taxon** | **Comparison** | **Wilcoxon rank sum statistic** | **p-value** |
|---|---|---|---|---|---|---|
| *P. glaber* | *P. cardinalis* | *P. eatonii* | *P. laevis* | H1H2 vs. H1B2 | 8973 | 0.3772 |
| *P. glaber* | *P. cardinalis* | *P. eatonii* | *P. laevis* | H1H2 vs. H2B1 | 14994 | 0.1215 |
| *P. glaber* | *P. cardinalis* | *P. eatonii* | *P. laevis* | H1H2 vs. H1B1 | 299681 | 4.273 e -11 |
| *P. glaber* | *P. cardinalis* | *P. eatonii* | *P. laevis* | H1H2 vs. H2B2 | 685778 | < 2.2 e -16 |
| *P. glaber* | *P. cardinalis* | *P. barbatus* | *P. virgatus* | H1H2 vs. H1B2 | 48222 | 0.0005119 |
| *P. glaber* | *P. cardinalis* | *P. barbatus* | *P. virgatus* | H1H2 vs. H2B1 | 93241 | 0.7091 |
| *P. glaber* | *P. cardinalis* | *P. barbatus* | *P. virgatus* | H1H2 vs. H1B1 | 699720 | 2.23 e -8 |
| *P. glaber* | *P. cardinalis* | *P. barbatus* | *P. virgatus* | H1H2 vs. H2B2 | 1477880 | < 2.2 e -16 |
| *P. glaber* | *P. cardinalis* | *P. labrosus* | *P.putus/deaveri* | H1H2 vs. H1B2 | 15374 | 0.9145 |
| *P. glaber* | *P. cardinalis* | *P. labrosus* | *P.putus/deaveri* | H1H2 vs. H2B1 | 18553 | 0.4028 |
| *P. glaber* | *P. cardinalis* | *P. labrosus* | *P.putus/deaveri* | H1H2 vs. H1B1 | 415638 | 6.818 e -12 |
| *P. glaber* | *P. cardinalis* | *P. labrosus* | *P.putus/deaveri* | H1H2 vs. H2B2 | 908752 | < 2.2 e -16 |
| *P. laevis* | *P. eatonii* | *P. barbatus* | *P. virgatus* | H1H2 vs. H1B2 | 24168 | 0.361 |
| *P. laevis* | *P. eatonii* | *P. barbatus* | *P. virgatus* | H1H2 vs. H2B1 | 18315 | 0.09904 |
| *P. laevis* | *P. eatonii* | *P. barbatus* | *P. virgatus* | H1H2 vs. H1B1 | 916161 | < 2.2 e -16 |
| *P. laevis* | *P. eatonii* | *P. barbatus* | *P. virgatus* | H1H2 vs. H2B2 | 646253 | < 2.2 e -16 |
| *P. laevis* | *P. eatonii* | *P. labrosus* | *P.putus/deaveri* | H1H2 vs. H1B2 | 24843 | 0.01205 |
| *P. laevis* | *P. eatonii* | *P. labrosus* | *P.putus/deaveri* | H1H2 vs. H2B1 | 28031 | 0.9198 |
| *P. laevis* | *P. eatonii* | *P. labrosus* | *P.putus/deaveri* | H1H2 vs. H1B1 | 985176 | < 2.2 e -16 |
| *P. laevis* | *P. eatonii* | *P. labrosus* | *P.putus/deaveri* | H1H2 vs. H2B2 | 812050 | < 2.2 e -16 |
| *P.virgatus* | *P. barbatus* | *P. labrosus* | *P.putus/deaveri* | H1H2 vs. H1B2 | 92016 | 0.0002475 |
| *P.virgatus* | *P. barbatus* | *P. labrosus* | *P.putus/deaveri* | H1H2 vs. H2B1 | 139307 | 0.002765 |
| *P.virgatus* | *P. barbatus* | *P. labrosus* | *P.putus/deaveri* | H1H2 vs. H1B1 | 1014506 | < 2.2 e -16 |
| *P.virgatus* | *P. barbatus* | *P. labrosus* | *P.putus/deaveri* | H1H2 vs. H2B2 | 1291339 | < 2.2 e -16 |

Table S7. Phylogenetic correlations among traits in bee syndrome species estimated in *caper*, using the *P. barbatus* reference genome.

| Model | Correlation coefficient | p-value |
|---|---|---|
| Long stamen length, short stamen length | 0.9681 | 7.002 e -13 |
| Long stamen length, style length | 0.8690 | 3.212 e -07 |
| Long stamen length, tube length | 0.8600 | 5.816 e -07 |
| Long stamen length, tube width | 0.2061 | 0.3701 |
| Long stamen length, log nectar volume | 0.524 | 0.0148 |
| Long stamen length, long nectary area | 0.4602 | 0.0358 |
| Short stamen length, style length | 0.8505 | 1.0429 e -06 |
| Short stamen length, tube length | 0.8616 | 5.225 e -07 |
| Short stamen length, tube width | 0.2953 | 0.1937 |
| Short stamen length, log nectar volume | 0.5642 | 0.0077 |
| Short stamen length, log nectary area | 0.4239 | 0.0558 |
| Style length, tube length | 0.8147 | 6.921 e -06 |
| Style length, tube width | 0.2372 | 0.3006 |
| Style length, log nectar volume | 0.6115 | 0.0032 |
| Style length, log nectary area | 0.3502 | 0.1197 |
| Tube length, tube width | 0.3167 | 0.1619 |
| Tube length, log nectar volume | 0.5713 | 0.0068 |
| Tube length, log nectary area | 0.3915 | 0.0793 |
| Tube width, log nectar volume | 0.4180 | 0.0593 |
| Tube width, log nectary area | 0.3575 | 0.1116 |
| Log nectar volume, log nectary area | 0.5435 | 0.0109 |

Table S8. Results of phylogenetic ANOVA models implemented in *caper* testing the effect of pollination syndrome on floral trait variation, using the *P. barbatus* reference genome. Lambda refers to the estimated Pagel's lambda parameter.

| **Model** | **lambda** | **DF** | **F-value** | **p-value** |
|---|---|---|---|---|
| long stamen length ~ syndrome | 0 | 28 | 53.949 | 5.353 e -08 |
| short stamen length ~ syndrome | 0.55 | 28 | 65.668 | 8.009 e -09 |
| style length ~ syndrome | 0.49 | 28 | 44.413 | 3.124 e -07 |
| tube length ~ syndrome | 0 | 28 | 59.01 | 2.282 e -08 |
| tube width ~ syndrome | 0.69 | 28 | 48.652 | 1.387 e -07 |
| log nectar volume ~ syndrome | 0 | 28 | 31.35 | 5.411 e -06 |
| log nectary area ~ syndrome | 1 | 28 | 57.963 | 2.711 e -08 |
| length / width ratio ~ syndrome | 0.15 | 28 | 301.73 | < 2.2 e -16 |
| anthocyanin ~ syndrome | 1 | 28 | 617.75 | < 2.2 e -16 |
| PC1 ~ syndrome | 0.44 | 28 | 141.68 | 1.804 e -12 |
| PC2 ~ syndrome | 0.43 | 28 | 3.0295 | 0.09275 |
| adjusted hue ~ syndrome | 0.94 | 23 | 302.18 | 1.003 e -14 |
| chroma ~ syndrome | 0 | 23 | 156.68 | 9.475 e -12 |
| brightness ~ syndrome | 0 | 23 | 15.032 | 0.000763 |
| chromatic contrast bee ~ syndrome | 0 | 23 | 57.507 | 1.061 e -07 |
| chromatic contrast bird ~ syndrome | 0 | 23 | 112.83 | 2.42 e -10 |

Table S9. Results of phylogenetic ANOVA models implemented in *caper* testing whether floral traits in bee syndrome species differ according to whether or not the accession is a sister taxon to hummingbird syndrome species, using the *P. barbatus* reference genome. Lambda refers to the estimated Pagel's lambda parameter.

| **Model** | **DF** | **lambda** | **F-value** | **p-value** |
|---|---|---|---|---|
| long stamen length ~ sister status | 19 | 1 | 2.3794 | 0.1394 |
| short stamen length ~ sister status | 19 | 1 | 3.8259 | 0.0653 |
| style length ~ sister status | 19 | 1 | 2.5132 | 0.1294 |
| tube length ~ sister status | 19 | 0 | 0.4426 | 0.5139 |
| tube width ~ sister status | 19 | 1 | 0.9375 | 0.3451 |
| log nectar volume ~ sister status | 19 | 0 | 1.9712 | 0.1765 |
| log nectary area ~ sister status | 19 | 0 | 0.4395 | 0.5153 |
| length / width ratio ~ sister status | 19 | 0 | 1.7317 | 0.2038 |
| anthocyanin ~ sister status | 19 | 1 | 2.5258 | 0.1285 |
| PC1 ~ sister status | 19 | 0.7 | 2.6172 | 0.1222 |
| PC2 ~ sister status | 19 | 0 | 2.4233 | 0.136 |

Table S10. Statistical comparisons of RND distributions between taxa in a 5-taxon testing framework. Reference genome used for analyses was *P. barbatus*.

| B1 taxon | H1 taxon | H2 taxon | B2 taxon | Comparison | Wilcoxon rank sum statistic | p-value |
|---|---|---|---|---|---|---|
| *P. glaber* | *P. cardinalis* | *P. eatonii* | *P. laevis* | H1H2 vs. H1B2 | 12522 | 0.7515 |
| *P. glaber* | *P. cardinalis* | *P. eatonii* | *P. laevis* | H1H2 vs. H2B1 | 23602 | 0.3671 |
| *P. glaber* | *P. cardinalis* | *P. eatonii* | *P. laevis* | H1H2 vs. H1B1 | 451140 | 6.43 e -15 |
| *P. glaber* | *P. cardinalis* | *P. eatonii* | *P. laevis* | H1H2 vs. H2B2 | 1024847 | < 2.2 e -16 |
| *P. glaber* | *P. cardinalis* | *P. barbatus* | *P. virgatus* | H1H2 vs. H1B2 | 72199 | 0.0006686 |
| *P. glaber* | *P. cardinalis* | *P. barbatus* | *P. virgatus* | H1H2 vs. H2B1 | 139390 | .2658 |
| *P. glaber* | *P. cardinalis* | *P. barbatus* | *P. virgatus* | H1H2 vs. H1B1 | 1197275 | 4.56 e -14 |
| *P. glaber* | *P. cardinalis* | *P. barbatus* | *P. virgatus* | H1H2 vs. H2B2 | 2699681 | < 2.2 e -16 |
| *P. glaber* | *P. cardinalis* | *P. labrosus* | *P.putus/deaveri* | H1H2 vs. H1B2 | 21447 | 0.8419 |
| *P. glaber* | *P. cardinalis* | *P. labrosus* | *P.putus/deaveri* | H1H2 vs. H2B1 | 25371 | 0.6417 |
| *P. glaber* | *P. cardinalis* | *P. labrosus* | *P.putus/deaveri* | H1H2 vs. H1B1 | 415638 | 6.818 e -12 |
| *P. glaber* | *P. cardinalis* | *P. labrosus* | *P.putus/deaveri* | H1H2 vs. H2B2 | 908752 | < 2.2 e -16 |
| *P. laevis* | *P. eatonii* | *P. barbatus* | *P. virgatus* | H1H2 vs. H1B2 | 41996 | 0.08567 |
| *P. laevis* | *P. eatonii* | *P. barbatus* | *P. virgatus* | H1H2 vs. H2B1 | 27932 | 0.08469 |
| *P. laevis* | *P. eatonii* | *P. barbatus* | *P. virgatus* | H1H2 vs. H1B1 | 1471736 | < 2.2 e -16 |
| *P. laevis* | *P. eatonii* | *P. barbatus* | *P. virgatus* | H1H2 vs. H2B2 | 1126924 | < 2.2 e -16 |
| *P. laevis* | *P. eatonii* | *P. labrosus* | *P.putus/deaveri* | H1H2 vs. H1B2 | 40892 | 0.1303 |
| *P. laevis* | *P. eatonii* | *P. labrosus* | *P.putus/deaveri* | H1H2 vs. H2B1 | 53938 | 0.1573 |
| *P. laevis* | *P. eatonii* | *P. labrosus* | *P.putus/deaveri* | H1H2 vs. H1B1 | 1694462 | < 2.2 e -16 |
| *P. laevis* | *P. eatonii* | *P. labrosus* | *P.putus/deaveri* | H1H2 vs. H2B2 | 1455412 | < 2.2 e -16 |
| *P.virgatus* | *P. barbatus* | *P. labrosus* | *P.putus/deaveri* | H1H2 vs. H1B2 | 163296 | 6.843 e -07 |
| *P.virgatus* | *P. barbatus* | *P. labrosus* | *P.putus/deaveri* | H1H2 vs. H2B1 | 232637 | 5.015 e -06 |
| *P.virgatus* | *P. barbatus* | *P. labrosus* | *P.putus/deaveri* | H1H2 vs. H1B1 | 1707757 | < 2.2 e -16 |
| *P.virgatus* | *P. barbatus* | *P. labrosus* | *P.putus/deaveri* | H1H2 vs. H2B2 | 2121034 | < 2.2 e -16 |

#### Supplemental References


Andrews, S., Krueger, F., Segonds-Pichon, A., Biggins, L., Krueger, C., & Wingett, S. (2010). FastQC. *A quality control tool for high throughput sequence data*, *370*.

Chen, S., Zhou, Y., Chen, Y., & Gu, J. (2018). fastp: an ultra-fast all-in-one FASTQ preprocessor. *Bioinformatics*, *34*(17), i884-i890.

Danecek, P., Auton, A., Abecasis, G., Albers, C. A., Banks, E., DePristo, M. A., Handsaker, R. E., Lunter, G., Marth, G. T., & Sherry, S. T. (2011). The variant call format and VCFtools. *Bioinformatics*, *27*(15), 2156-2158.

Harborne, J. (1984). *Phytochemical methods 2nd ed*. Chapman and Hall.

Jarvis, D. E., Stevens, M. R., Carter, P., Lin, Y. F., Jaggi, K. E., Jijon, G., Kalt, T., Calixto, J., Standring, S., & Torres, K. (2025). Whole-genome assembly and annotation of the firecracker penstemon (Penstemon eatonii). *Journal of Heredity*, *116*(3), 373-381.

Jun, G., Wing, M. K., Abecasis, G. R., & Kang, H. M. (2015). An efficient and scalable analysis framework for variant extraction and refinement from population-scale DNA sequence data. *Genome research*, *25*(6), 918-925.

Li, H. (2011). A statistical framework for SNP calling, mutation discovery, association mapping and population genetical parameter estimation from sequencing data. *Bioinformatics*, *27*(21), 2987-2993.

Li, H. (2013). Aligning sequence reads, clone sequences and assembly contigs with BWA-MEM. *arXiv preprint arXiv:1303.3997.*

Li, H., Handsaker, B., Wysoker, A., Fennell, T., Ruan, J., Homer, N., Marth, G., Abecasis, G., & Durbin, R. (2009). The sequence alignment/map format and SAMtools. *Bioinformatics*, *25*(16), 2078-2079.

Maia, R., Gruson, H., Endler, J. A., & White, T. E. (2019). pavo 2: new tools for the spectral and spatial analysis of colour in R. *Methods in Ecology and Evolution*, *10*(7), 1097-1107.

Minh, B. Q., Hahn, M. W., & Lanfear, R. (2020). New methods to calculate concordance factors for phylogenomic datasets. *Molecular biology and evolution*, *37*(9), 2727-2733.

Nguyen, L.-T., Schmidt, H. A., Von Haeseler, A., & Minh, B. Q. (2015). IQ-TREE: a fast and effective stochastic algorithm for estimating maximum-likelihood phylogenies. *Molecular biology and evolution*, *32*(1), 268-274.

Schindelin, J., Arganda-Carreras, I., Frise, E., Kaynig, V., Longair, M., Pietzsch, T., Preibisch, S., Rueden, C., Saalfeld, S., & Schmid, B. (2012). Fiji: an open-source platform for biological-image analysis. *Nature methods*, *9*(7), 676-682.

Smith, S. A., & O'Meara, B. C. (2012). treePL: divergence time estimation using penalized likelihood for large phylogenies. *Bioinformatics*, *28*(20), 2689-2690.

Smith, S. D. (2014). Quantifying color variation: improved formulas for calculating hue with segment classification. *Applications in Plant Sciences*, *2*(3), 1300088.

Stevens, J. T., Wheeler, L. C., Williams, N. H., Norton, A. M., & Wessinger, C. A. (2023). Predictive links between petal color and pigment quantities in natural Penstemon hybrids. *Integrative and Comparative Biology*, *63*(6), 1340-1351.

Stone, B. W., & Wessinger, C. A. (2024). Ecological diversification in an adaptive radiation of plants: the role of de novo mutation and introgression. *Molecular biology and evolution*, *41*(1), msae007.

Vorobyev, M., & Osorio, D. (1998). Receptor noise as a determinant of colour thresholds. *Proceedings of the Royal Society of London. Series B: Biological Sciences*, *265*(1394), 351-358.

Wessinger, C. A., & Rausher, M. D. (2015). Ecological transition predictably associated with gene degeneration. *Molecular biology and evolution*, *32*(2), 347-354.

Zhang, C., Rabiee, M., Sayyari, E., & Mirarab, S. (2018). ASTRAL-III: polynomial time species tree reconstruction from partially resolved gene trees. *BMC bioinformatics*, *19*(6), 153.
